# Biophysical limits of ultrafast cellular motility

**DOI:** 10.1101/2024.08.22.609204

**Authors:** Ray Chang, Manu Prakash

## Abstract

Many single-celled organisms and specialized cell types can surprisingly achieve speed and acceleration significantly faster than multicellular counterparts. These remarkable cellular machines must integrate energy storage and amplification in actuation, latches for triggered release, and energy dissipation without failure — all implemented in macro-molecular assemblies inside a single cell. However, a universal biophysical framework that can comparatively evaluate extreme cellular motility remains lacking. Scaling laws have long been recognized as powerful tools for revealing universal principles in physical systems. We map the atlas of ultrafast motility for single cells across the tree of life. We then introduce a new quantitative framework that can be used to evaluate and compare extreme acceleration, speed, area strain rate, volume expansion strain rate, and density changes in single cells. Recognizing that many single cells operate in low-Reynolds number environments, we introduce a new dimensionless number, the “cellular acceleration number,” based on energy dissipation at this scale. Using this new framework, we discover a scaling law between the cellular acceleration number and the transient Reynolds number, valid across six orders of magnitude in a range of single-cell organisms. We further generalize these ideas by placing various trigger, actuation, and dissipation mechanisms within the same framework and estimating the fundamental limits of speed and acceleration at the cellular scale. We conclude with a detailed summary of the range of functions implemented via ultrafast cellular phenomena, laying down a quantitative foundation for extreme biophysics at the cellular scale.

## Introduction

As Sun Tzu wrote in the book ‘The Art of War,’ ‘Speed is the essence of war. Exploit the enemy’s unpreparedness; attack him unawares; take an unexpected route.’^1^ This is equally true for both multicellular organisms and single cell organisms, as we are all constantly “at war” against various evolutionary pressures. Indeed, many interesting biological phenomena were discovered when we examine the physical extremes of biological processes. For example, the cheetah is the fastest running land animal, reaching a horizontal speed of 25 m/sec and an acceleration of 1.5*g*^2^. A grasshopper can jump can reach an acceleration of 18*g*^3^. Plants have also evolved various ultrafast mechanisms for nutrient acquisition and reproduction, exhibiting velocities and accelerations comparable to those of animals. The suction trap of bladderworts, for instance, can achieve a fluid velocity of 1.5 m/sec and an acceleration of 600*g*, trapping water fleas and worms for their food^4^. The flower of bunchberry has a catapult mechanism, ejecting its pollen at a speed of 7.5 m/sec and an acceleration of 2,400*g*, as its pollination is self-incompatible and requires this ultrafast mechanism to enhance insect pollination and potentially wind pollination^5^. On an even faster acceleration scale, the famous mantis shrimp appendage strike can reach a velocity of 31 m/sec and an acceleration of 25,000*g*^6^, while the mandible strike of trap-jaw ants can reach a velocity of 64 m/sec and an acceleration of 100,000*g*^7^, used for both attack and escape jump^8^. These examples from the multicellular world, though impressive, require specialized tissues to achieve these ultrafast movements. However, none surpasses the 5,400,000*g* acceleration of nematocyst discharge in *Hydra*, which ultimately is a single-cell phenomenon performed by specialized cells called nematocytes^9^. Despite past fascination with many macroscopic examples, the extremes in cellular world remain largely unexplored, offering a rich field for discovery.

Ultrafast phenomena pose specific challenges at the single-cell level. For example, all the energy required for the ultrafast mechanisms has to be contained within a single cell’s size, only to be deployed when the proper stimuli are present. For the nematocytes of *Hydra*, the task is somewhat easier as their discharge is a single-shot event, but these specialized cells are supported by surrounding tissues in the organism. However, single-cell organisms with ultrafast motility lack this luxury, as they must also perform all the necessary functions as a single unit. Aside from that, they need to ensure to remain viable even after the deployment of their ultrafast mechanisms without any support from connective tissues, as extreme velocity and acceleration usually indicate the presence of extreme shear and forces. These challenges pose additional evolutionary pressures on organisms capable of ultrafast motility.

In this work, we first create a map of single-cell organisms that demonstrate ultrafast motility, including locomotion, contractility, volume expansion, and density changes. Next, we classify and quantitatively compare these behavioral phenotypes. In order to understand ultrafast acceleration in a cellular context, we propose a new non-dimensional number, the “cellular acceleration number” (cAc), which is the ratio of relaxation time scale in Stokes flow to the acceleration time scale. This approach enables us to determine whether a phenomenon at microscopic scales qualifies as ultrafast acceleration. Using this new quantitative framework and the atlas described earlier, we discover a near-linear scaling between the cellular acceleration number and the transient Reynolds number for ultrafast cellular phenomena across six orders of magnitude. The slope of this curve allows us to estimate the energy conversion efficiency of ultrafast motility in dissipative media. This energy conversion efficiency, together with energy mass density and energy release time scale, can be used to estimate the fundamental speed and acceleration limits of multiple ultrafast mechanisms at the microscale. We conclude with a comprehensive review of the functions of cellular ultrafast phenomena and their corresponding evolutionary adaptations.

### Cellular Olympics: Classification of behavioral phenotypes in ultrafast motility

Ultrafast motility evolved in single-cell organisms from a wide range of phylogenetic origins, displaying diverse morphology, sizes, sub-cellular organization, and behavioral phenotypes (Fig. 1). Single cells exhibiting ultrafast motility vary dramatically in size, spanning at least three orders of magnitude, and can be found in bacteria, archaea, and various major groups of eukaryotes. In the following sections, we classify different behavioral phenotypes in ultrafast motility and later compare them quantitatively. We collected these data from published works and re-analyzing the figures or videos from those works, which together gave us 17 acceleration, 224 speed, 38 cilia morphology, 5 area strain rate, 9 volume expansion strain rate, and 4 density change rate data. Note that this classification is not phylogenetic but a convenient way to comprehensively cover the ultrafast cellular phenomena reported so far.

**Figure 1.**
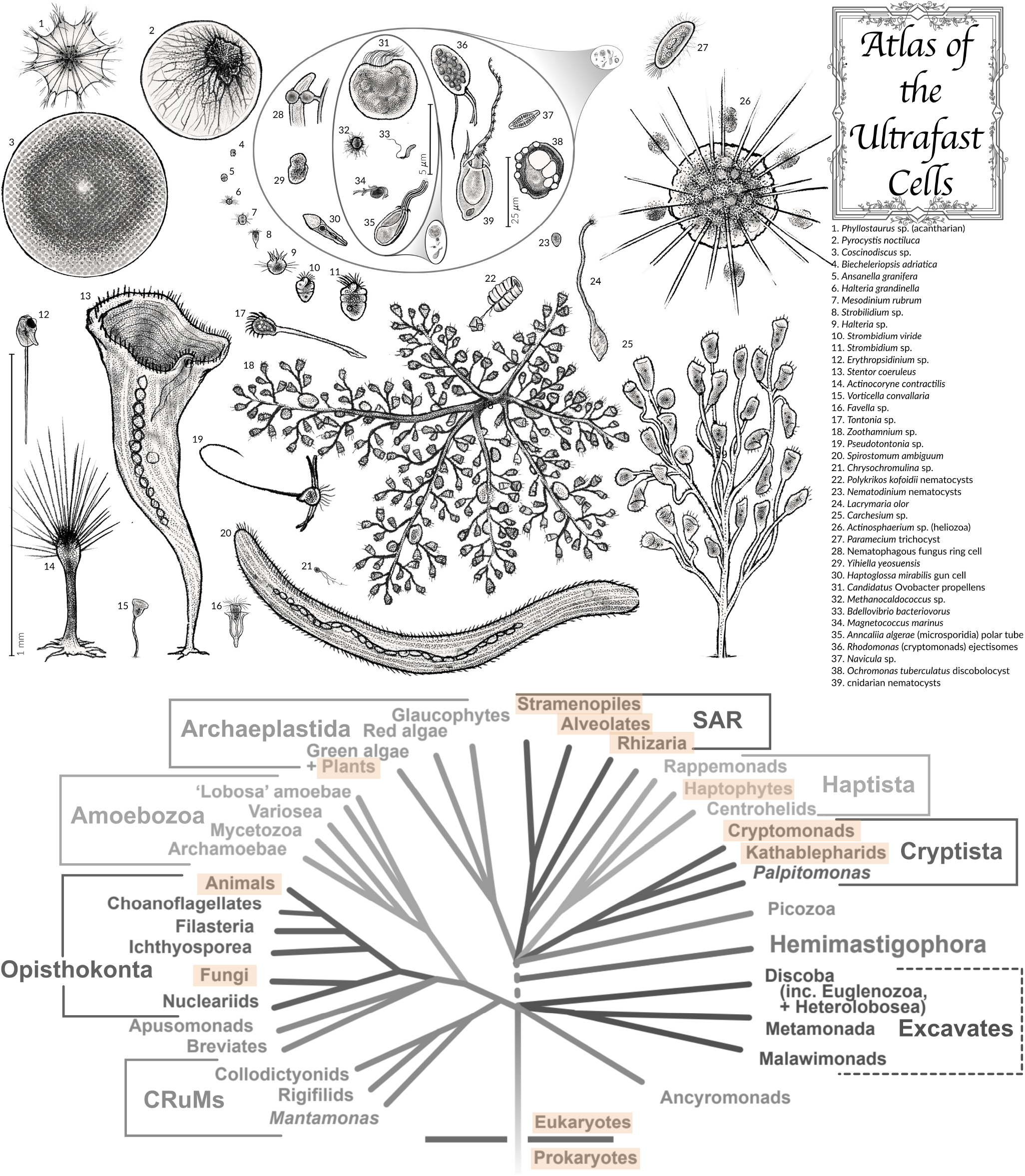
Organisms with ultrafast cellular motility shown in relative scale to each other and their presence highlighted in orange on the eukaryotic phylogenetic tree. Ultrafast organisms, which come in a wide range of sizes, shapes, and behavioral phenotypes, are present in multiple major branches of the eukaryotic phylogenetic tree. This list includes fast translocation, contraction, extrusion of organelles, and even fast volume and area changes. The phylogenetic tree of eukaryotes is adapted from the work by Alastair Simpson^10^.

#### Hydrodynamic swimmers

We first classify organisms whose motility is characterized by a smooth swimming pattern through thrust generated from cilia or flagella as “hydrodynamic swimmers”. Among them, the fastest swimming bacteria known to date is *Candidatus* Ovobacter propellens (Fig. 1, #**31**)^11^. These large bacteria (∼ 5*μ*m in size) have a huge flagellar tuft consisting of about 400 flagella. They prefer an oxygen tension of about 0.5% atmospheric saturation in marine sediments, where the oxygen gradient is usually steep. They swim continuously at a speed of 0.6-0.7 mm/sec, with some reaching up to 1 mm/sec (200 BL/sec). However, if we rank bacteria according to their maximum normalized speed, the title goes to *Magnetococcus marinus* (MC-1), which can swim at 0.5 mm/sec despite being only 1 *μ*m in size (Fig. 1, #**34**)^12^. MC-1 carries a magnetosome chain that allows it to sense the magnetic fields, and it also prefers microoxic conditions. MC-1 has two flagella bundles at each pole of the cell, which work cooperatively to push on one end and pull on the other, enabling their fast swimming speed and quick reorientation. Aside from bacteria, archaea such as *Methanocaldococcus villosus* and *Methanocaldococcus jannaschii* (Fig. 1, #**32**) can also reach swimming speed of up to 468 BL/sec and 393 BL/sec, respectively^13^. They are thermophilic hydrogenotrophic methanogens, preferring high temperatures near hydrothermal vents, and live on hydrogen and carbon dioxide, producing methane as a byproduct. Their swimming patterns are characterized by fast, straight swimming to relocate, and slower zigzag swimming trajectories when near a surface^13^.

Another example of fast bacteria is *Bdellovibrio bacteriovorus* (Fig. 1, #**33**), a predatory bacterium with a single flagellum capable of penetrating and replicating within bacterial prey^14^. For strain HD100, this 1-*μ*m-large bacterium can swim up to 160 *μ*m/sec. Early hypotheses suggested that they use their fast motility to drill and penetrate the prey cell, but later genetic knockout studies showed that flagellar motility is not essential for prey entry and thus only plays a role in efficient prey encounter^14^.

#### Hydrodynamic jumpers

We next denote organisms whose swimming trajectories are characterized by slower regular swimming mode interspersed with sudden high speed “jumps” as “hydrodynamic jumpers.” Many small-sized ciliates have evolved the ability to perform sudden jumps, and they are collectively referred to as “jumping ciliates” (see Tamar 1979 for a comprehensive review^15^). They include *Halteria* (Fig. 1, #**6**&#**9**), *Mesodinium* (Fig. 1, #**7**), *Askenasia, Cyclidium, Pleuronema, Strobilidium* (Fig. 1, #**8**), *Urotricha, Uronychia, Dinophrys*, and some weaker jumpers like *Strombidium* (Fig. 1, #**10**&#**11**) and *Stylonychia*. Take *Halteria grandinella* as an example: this 20-*μ*m-large oligotrich normally swims forward at a speed of 0.2-0.3 mm/sec (10-15 BL/sec), but when disturbed by flow or a high concentration of KCl, it performs a powerful backward jump at a speed of 3.2 mm/sec (160 BL/sec). The maximum normalized speed of jumping ciliates typically fall between 50-200 BL/sec, with *Strobilidium* sp. having the fastest recorded speed of 2.76 cm/sec for a 76-*μ*m cell (361 BL/sec)^16^. These fast jumps are enabled by their relatively long cilia and an exceptionally high jump in cilia beating frequency from 105 Hz to 260 Hz^17^ (see Section “Actuation” below for a full discussion).

Aside from jumping ciliates, many dinoflagellates also are capable of hydrodynamic jumps. Examples include *Ansanella granifera* (Fig. 1, #**5**)^18^, *Yihiella yeosuensis* (Fig. 1, #**29**)^19^, and *Biecheleriopsis adriatica* (Fig. 1, #**4**)^20^, *Heterocapsa rotundata*^21^ and *Gymnodinium simplex*^21^. The champion of fastest swimming speed is *Ansanella granifera*, with a size of 10.5 *μ*m capable of swimming at 1603 *μ*m/sec (153 BL/sec)^18^, while the fastest normalized swimming speed goes to *Yihiella yeosuensis*, with a size of 8 *μ*m capable of swimming at 1572 *μ*m/sec (197 BL/sec). Functionally, while these jumps have an obvious role in predator escape, they can also be used for ambush attacks^22^ or to escape from their own nutrient-depleted boundary layers to enhance the diffusion of nutrients.^23^ It is also believed that their fast swimming speed can help them reach deep-water nutrients.^19^

In addition to jumping ciliates and dinoflagellates, sudden jumps are also reported in *Bodo saltans*^24^, a free-living kinetoplastid, and *Resultomonas moestrupii* (formerly *Pedinomonas mikron*)^25^, a primitive green algae. However, no swimming or jumping speeds have been reported for them. It is also important to note that although cryptophyte species are capable of jumping, their jumps are mediated by the ejection of ejectisomes and are thus categorized as ‘cell protrusions or extrusomes’^26^.

#### Elastic jumpers

Some organisms, instead of jumping through fast ciliary or flagellar beating, have a long appendage that is normally stretched out and can jump by contracting or recoiling their long appendages back to their cell body. Confirmed examples from ciliates include *Tontonia*^16^ (Fig. 1, #**17**), *Pseudotontonia*^16^ (Fig. 1, #**19**), and *Spirotontonia*^27^. When their appendages contract, the cell bodies of *Pseudotontonia* and *Tontonia* can achieve backward jumps at 653 BL/sec and 1470 BL/sec, respectively.^16^ Preliminary electron microscopy studies on the tail of *Tontonia appendiculariformis* showed that the tail is composed of numerous nearly parallel membranous tubes and septa connected to small bundles of microfilaments, and the tubular layers are separated by hyaloplasm containing many mitochondria.^28^ A similar mechanism has also evolved in haptophytes such as *Chrysochromulina* (Fig. 1, #**21**)^29^. When the microtubule array in the haptonema (the unique appendage present only in haptophytes) switches its conformation - a calcium driven process presumably involving Ca^2+^-binding microtubule-associated proteins - the haptonema coils back and causes a rapid jump of the cell body at a speed of 6 mm/sec (or 600 BL/sec).^29^ Another ciliate, *Zoothamnium pelagicum* (Fig. 1, #**18**), while not having a contractile tail, can contract the entire pseudocolony and achieve a medusa-like locomotion at a speed of 110 mm/sec (∼ 250 pseudocolony size/sec).

#### Surface-based motility

We next designate organisms whose motility mechanisms depend on the presence of a solid substrate for “surface-based motility.” One example of this is diatoms. Diatoms (Fig. 1, #**37**), despite lacking cilia or flagella (except in male gametes of some species), are capable of gliding on surfaces through actomyosin motility system linked to secreted adhesive polysaccharide^30^. Although the average gliding speed on the surface is usually low (∼ 2-20 *μ*m/sec, *<*1-5 body lengths (BL)/sec)^31^, high-speed imaging has revealed that the gliding of *Navicula* and *Nitzschia* species is characterized by jerky movements, with bursts of instantaneous speed up to 250 *μ*m/sec for the *Navicula* diatom, corresponding to 125 BL/sec^30^. A recent study suggested that this jerky motion might result from the misalignment in direction between the extruded polysaccharide and the direction of cell motion, but more studies need to be done.^31^

#### Contraction without translocation

We next consider examples where the contraction of either the entire cell body or their appendages is not mainly used for translocation. These organisms are either attached to a substrate with a stalk, or their motility is symmetric and thus only result in little net translocation after a complete actuation and relaxation cycle. Examples from ciliates include contractile heterotriches like *Spirostomum*^32^ (Fig. 1, #**20**) and *Stentor*^33^ (Fig. 1, #**13**), contractile sessilids like *Vorticella* (Fig. 1, #**15**), *Carchesium* (Fig. 1, #**25**), and *Zoothamnium* (Fig. 1, #**18**) (see Ryu et al. 2022 for a review on the biomechanics of various contractile sessilids species^34^), and tintinnids like *Favella* (Fig. 1, #**16**)^35^. Take *Spirostomum* and *Stentor* as examples, they are both giant single-cell ciliates that can grow to mm-scale and can contract their cortical myoneme to reduce their body length to less than 50% within 5 msec. For *Spirostomum ambiguum* in particular, this corresponds to a contraction speed of 0.2 m/sec and an acceleration of 15*g*^32^.

Fast contraction has also been observed in various heliozoan species (see Davis 1994 for a comprehensive review of the biophysics of this process^36^). For species like *Echinosphaerium, Actinophaerium* (Fig. 1, #**26**), *Heterophrys* and *Actinophrys*, their outstretched axopodia can rapidly contract when in contact with food organisms, bringing the prey near the body surface^37^. For *Echinosphaerium*, the contraction speed of axopodia is estimated to be more than 2 mm/sec^37^. One special benthic heliozoan species, *Actinocoryne contractilis* (Fig. 1, #**14**), can not only contract their axopodia but can also contract their stalk. The stalk, 300-400 *μ*m long when fully stretched, can be shortened by 90% within 50-60 msec, causing a speed up to 10 cm/sec (or 600 BL/sec) and an acceleration of 10*g*^38^.

Similar behavior is also observed in dinoflagellate species *Erythropsidinium* (Fig. 1, #**12**) and *Greuetodinium*. They are both characterized by the presence of ocelloids and a fast contractile appendage commonly referred to as ‘piston’^39^. While the contraction speed of the piston has never been reported in *Greuetodinium* (which might just be the early division stage of *Erythropsidinium* rather than a different genus or species^39^), the contraction speed of the piston in *Erythropsidinium* can go up to 3.3 cm/sec (∼ 330 BL/sec). While *Erythropsidinium* mostly use their piston in static mode for feeding (i.e., no major translocation of cell body after a complete contraction/extension cycle), in locomotion mode they can cause the cell to move at a net speed of 1 mm/sec.^39^

#### Cell protrusions or extrusomes

Opposite to cell contractions, some organisms are also capable of extending part of the cell or have specialized organelles that protrudes or extrudes. Specialized exocytic organelles that can extrude materials at very high speed, which is commonly referred to as extrusomes (see the reviews by Hausmann 1978^40^ and Rosati & Modeo 2003^41^ for a comprehensive discussion on this topic). Examples of fast extrusomes include trichocysts from many ciliate species (with *Paramecium* being the most studied example (Fig. 1, #**27**)), toxicysts (including pexicysts) from many predatory ciliates like *Didinium, Lacrymaria*, and *Dileptus*, dinoflagellate nematocysts from *Polykrikos* (Fig. 1, #**22**) and *Nematodinium* (Fig. 1, #**23**), ejectisomes from cryptomonads (Fig. 1, #**36**) and kathablepharids, cnidarian nematocysts (also known as cnidocysts) from all cnidarian species (Fig. 1, #**39**), and discobolocyst from specific chrysophyte species such as *Ochromonas tuberculatus* (Fig. 1, #**38**), *Cyclonexis annularis*, and *Chromulina erkensis* (see Hibberd 1970 for the full list of species with discobolocysts^42^).

Among these, cnidarian nematocyst holds the record for the fastest experimentally proven speed (37 m/sec) and acceleration (5.4 × 10^6^*g*) at the single-cell level^9^. However, it is suspected that the true fastest record might belong to the discobolocysts of *Ochromonas tuberculatus*. It was calculated that the initial velocity of discobolocyst extrusion can reach up to 260 m/sec, based on the observed travel distance of the cell body after the explosion of discobolocyst, hydrodynamic drag, and momentum conservation^43^, although this calculation has never been validated with direct measurement of extrusion velocity. The trichocyst of paramecium can achieve a speed of 1 cm/sec given a 3 *μ*m organelle size^44^, while the ejectisome from cryptophyte species *Storeatula* can go up to 2 cm/sec given a 0.8 *μ*m organelle size.^26^ So far, no precise velocity measurements have been made for toxicysts or pexicysts.

Aside from extrusomes, two groups of parasites have evolved similar behavioral phenotypes by developing a cell protrusion that extrudes at high speed to deliver infectious cargo into the host. One of these groups is microsporidia, a group of obligatory intracellular parasitic fungi (Fig. 1, #**35**), which infect their hosts through an infective spore containing an extrusive organelle called a polar tube (some early literature also categorized the polar tube as an extrusome^40^, but this classification is less common in recent literature). When triggered, the polar tube of *Anncaliia algerae* can eject at a speed of 300 *μ*m/sec (or 100 BL/sec), penetrate the membrane of the host cell, and deliver the infectious cargo into the cell^45^. The other group is the gun cell of *Haptoglossa* (Fig. 1, #**30**), a group of parasitic oomycetes that infect nematodes or rotifers. The gun cell of *Haptoglossa mirabilis* is about 15-18 *μ*m long and features a complex ejection system composed of a bore, a plug, a projectile, and a flexible tube. When triggered, the ejection system everts and penetrates the cuticle of the host, completing the infection within 0.1 sec.^46^ No precise velocity measurements have been made for *Haptoglossa* so far.

Lastly, another example of cells with fast protrusion is *Lacrymaria olor*(Fig. 1, #**24**), a freshwater predatory ciliate characterized by a teardrop-shaped cell body (∼ 40*μ*m) connected to a “head” with a long slender neck. The neck is driven by the head cilia to search the surrounding space, extending and contracting at a speed of up to 1500 *μ*m/sec^47^. The combined effect of driving force from the head, hydrodynamic interaction, and neck elasticity gives rise to a programmable chaotic search behavior^48^. As the neck extends and contracts, the surface area changes at a rate corresponding to a 400%/sec area strain rate, much faster than the previously reported theoretical maximum area strain rate of 250%/sec based on actomyosin^49^. This remarkable speed is possible due to the sequential deployment from the curve-creased origami of stored cell membrane and microtubule sheets^50^.

#### Extreme volume and density changes

Finally, ultrafast processes at cellular scale are not necessarily limited to fast speed or acceleration; cells can also employ rapid changes in area, volume, or density to achieve various functions. For example, *Pyrocystis noctiluca* (Fig. 1, #**2**), a nonmotile dinoflagellate, can rapidly expand its volume by 5 times within 10-15 minutes after cell division to achieve positive buoyancy.^51^ Acantharian (Fig. 1, #**1**) can also rapidly expand their outer cytoplasm through contracting myonemes.^52^ Although it is commonly believed that this cytoplasm expansion is used to adjust drag and buoyancy, actual experimental evidence is limited. Large diatoms like *Coscinodiscus* (Fig. 1, #**3**) are also able to rapidly change their density at a rate of 584 kg/m^3^/sec^53^, presumably through rapid ionic exchanges^54^.

Aside from buoyancy adjustment, cells can also utilize rapid volume changes to attack prey. For example, the ring cells of nematophagous fungi (such as in *Arthrobotrys dactyloides*, formerly known as *Drechslerella dactyloides*) can triple in size within 0.1-1 second to capture nematodes, causing a volume expansion strain rate of 3000%/sec^55^ (Fig. 1, #**28**). Typical animal cell lines like CHO cells would die if the volume expansion strain rate caused by the water influx exceeds 0.5%/sec.^56^

### Quantitative framework for ultrafast cellular motility

So far, we have seen many examples of single cells using fast movements or rapid changes in surface area and volume to achieve various functions. These cells span across the tree of life, and as shown in Figure 1, range from 1-4 mm to 0.5 *μ*m. To systematically compare and contrast these phenomena and identify the most extreme cases worthy of the title ‘ultrafast,’ we develop a comprehensive quantitative framework to evaluate and rank these fast processes based on measurable parameters. By identifying outliers and extreme cases, we can uncover potentially new biological mechanisms and biophysical principles that are not yet fully understood, thus deepening our understanding of cellular biology in extremes.

#### Ultrafast acceleration

##### Extreme acceleration in low Reynolds number

We first start by examining accelerations at cellular scales. In our previous discussion, we mentioned several examples such as the 10^6^*g* acceleration of nematocyst discharge^9^ and the 15*g* acceleration of *Spirostomum* contraction^32^, and more examples are summarized in Figure S1. Past studies either directly compare the acceleration magnitude with multicellular examples (Fig. S1A), or they normalize the acceleration by body length to account for the wide size range differences (Fig. S1B). However, neither method is justifiable when considering the low-Reynolds-number physics. Unlike multicellular organisms, single-cell motility is always embedded in water, where the relative importance of viscous dissipation of water fundamentally changes with the length scale. One common way to evaluate this is by looking at the Reynolds number, a dimensionless number that quantifies the relative importance of inertia and viscous forces in fluid flow. As most cellular motility occurs in the low Reynolds number regime (Fig. 2B), inertia is often negligible at the cellular scale, and motions can abruptly stop with seemingly high deceleration.

**Figure 2.**
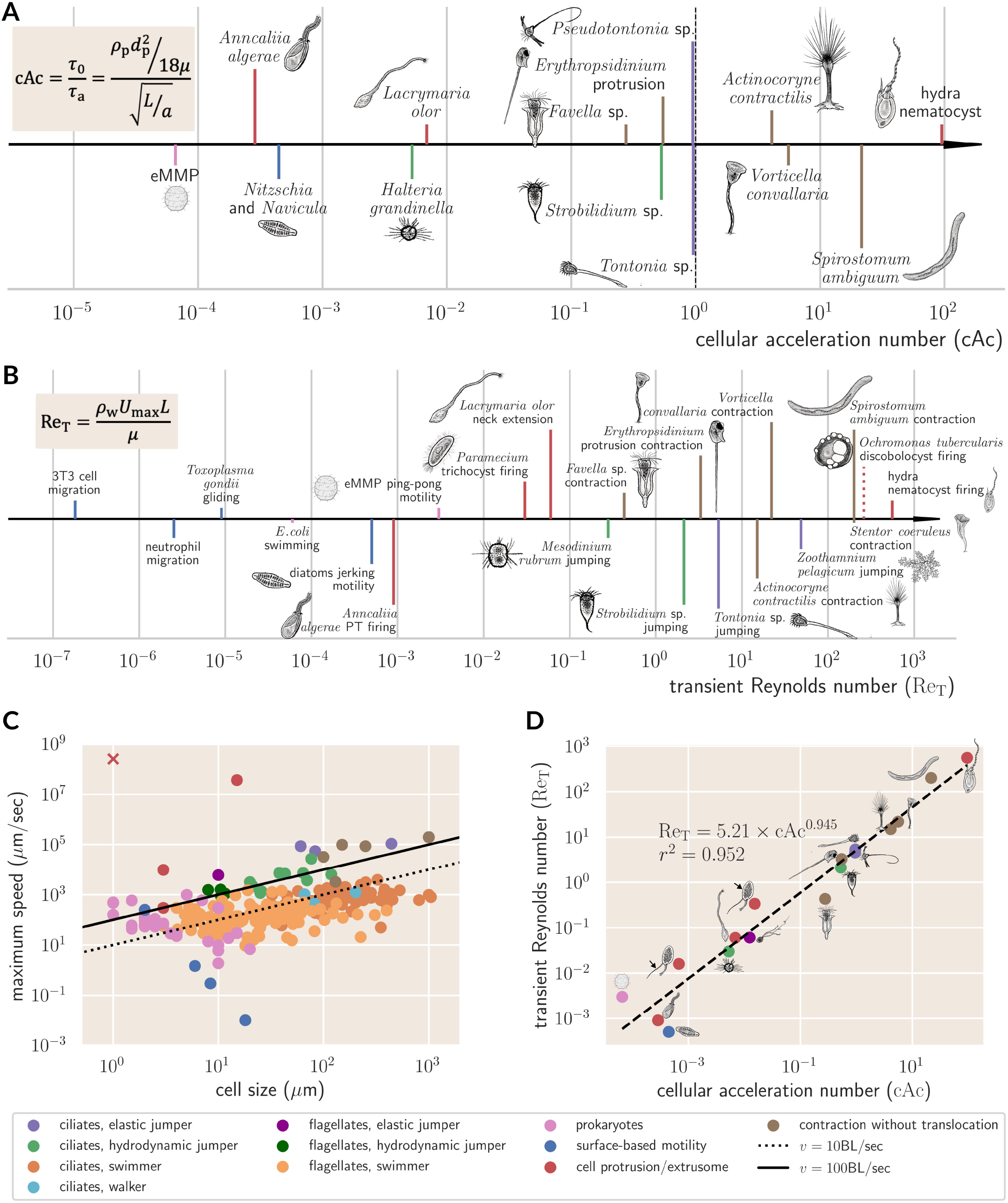
A novel scaling law for ultrafast cellular phenomena. (A) Fast acceleration at cellular scales. As single-cell organisms with ultrafast motility are embedded in a viscous, dissipative environment, we present a new dimensionless number to account for the effect of Stokes drag to compare the acceleration magnitude. We thus define the cellular acceleration number (cAc) as the ratio between the relaxation time scale in Stokes flow (*τ*_0_) and the acceleration time scale (*τ*_a_). When cAc is greater than 1, the accelerations we observe cannot be explained by the naive hydrodynamic effect of low Reynolds number flow and thus qualify as “ultrafast acceleration.” Different colors indicate various behavioral phenotypes (color code shown below). Examples qualifying as ultrafast acceleration include extrusomes (hydra nematocysts) and contractions without translocation (e.g., *Spirostomum ambiguum, Vorticella convallaria, Actinocoryne contractilis*). (B) Fast velocities at cellular scales. Because ultrafast organisms achieve maximum speed only transiently, we compare their speed using transient Reynolds number (Re_T_) defined by their maximum speed (*U*_max_). While many ultrafast velocities at the cellular scale are still in the low Reynolds number regime, many single-cell organisms achieve maximum velocities in the intermediate Reynolds number regimes (1 *<* Re_T_ *<* 1000). (C) Statistics of maximum speed at cellular scales. Most organisms can only achieve a maximum velocity at around 10 body lengths/sec, justifying the definition of a normalized speed greater than 100 body lengths/sec as ultrafast velocity. The ‘×’ symbol indicates the discobolocyst of *Ochromonas tuberculatus*, which is an inferred number that is not experimentally validated.^43^ (D) As acceleration and maximum velocity are both limited by energy sources and the viscous dissipation from the surrounding, we compare the two dimensionless numbers, Re_T_ and cAc. We discovered a near-linear scaling between transient Reynolds number and cellular acceleration number for ultrafast motility across multiple scales and various behavioral phenotypes. This suggests a constant energy conversion efficiency of 8% from acceleration to observed speed. (See main text for calculation details.)

For example, consider the Stokes drag deceleration on a spherical object moving at speed *v* with density *ρ*_p_ and diameter *d*_p_ in a fluid with dynamic viscosity *μ*. The expected deceleration from Stokes drag alone can be calculated as 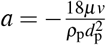. As we will mention later, for most organisms the maximum speed is about 10*d*_p_/sec^57^. Plugging in the relevant numbers, we can show that the naive hydrodynamic deceleration on an organism is 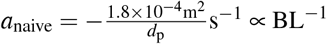. This means that for a bacterium with a size of 1.8 *μ*m, it does not require any special mechanism to achieve a 10*g* deceleration! Additionally, attempting to normalize acceleration with body length only exacerbates the issue, as *a*_naive_*/*BL ∝ BL^*−*2^. This motivates us to search for a new dimensionless number that can account for energy dissipation effects and properly compare accelerations at cellular scale.

##### Evaluating acceleration using cellular acceleration number

To account for the extreme deceleration or high acceleration just from Brownian motions for micron-scale object in water, we define a new dimensionless number based on the ratio between two time scales. One is the relaxation time scale of particles in Stokes flow, which is 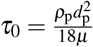. The other is the acceleration time scale, which can be defined by the characteristic acceleration (*a*) and characteristic length scale (*L*) of a biological process as 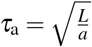. The ratio of these two time scales defines the cellular acceleration number as 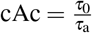. When the cellular acceleration number is less than one, the observed acceleration does not outperform the naive hydrodynamic effect of low Reynolds number flow and thus should not be regarded as ultrafast. Physically, the cellular acceleration number can also be regarded as a special case of Deborah number, with the characteristic time scale of acceleration replacing the time of observation in the common definition.

Take the contraction of *Spirostomum* as an example (Fig. 1, #**20**). As we mentioned earlier, *Spirostomum ambiguum* is a giant single-cell ciliate that can grow up to 1-4 mm, and it possesses an ability to contract itself to less than 50% body length in just 5 msec^32^. The contraction process is characterized by a rapid acceleration followed by rapid deceleration, both have a magnitude of 15*g*. We can use the cellular acceleration number to evaluate whether these accelerations and decelerations are greater than the naive hydrodynamic effects. Taking its length (∼ 1 mm) as the characteristics length scale for the calculation of both *τ*_0_ and *τ*_a_, its relaxation time in Stokes flow is about 56 msec and its acceleration time scale is about 2.6 msec. This gives a cellular acceleration number of 21.5, which means the acceleration and deceleration of *Spirostomum* contraction process cannot be ascribed to just simple hydrodynamics effects. This is consistent with our knowledge that the fast contraction of *Spirostomum* comes from the calcium-triggered release of energy stored in the giant spasmin binding protein (GSBP)-spasmin protein complex located at the cell cortex^58^, while the rapid deceleration comes from the strain-induced jamming transition of entangled intracellular organelles.^59^

We can next look at another example from microsporidia species *Anncaliia algerae* (Fig. 1, #**35**). The polar tube ejection from the 3 *μ*m-sized spore has an acceleration of 0.061 m/sec^245^. We can thus calculate *τ*_0_ and *τ*_a_ as 2 *μ*sec and 7 msec, respectively, and this yields a cellular acceleration number of 2.8 × 10^*−*4^. This indicates that the acceleration we observed in microsporidia polar tube ejection is much weaker compared to the acceleration you would expect just from naive hydrodynamic effects and thus should not be regarded as extreme. Interestingly, if you calculate the normalized acceleration for *Spirostomum ambiguum* and *Anncaliia algerae*, their *a/*BL are 1.5 × 10^5^ s^*−*2^ and 2.0 × 10^4^ s^*−*2^, respectively, which do not look that different. This again shows that a dimensionless measurement using cellular acceleration number is a much better approach than normalized acceleration to compare accelerations at cellular scale.

We can now compare the examples of fast acceleration shown in Figure S1 using cellular acceleration number (Fig. 2A). Lines of different colors indicate different behavioral phenotypes. The examples that qualify as ultrafast acceleration are either extrusomes (hydra nematocysts, red lines) or contraction without translocation (*Spirostomum ambiguum, Vorticella convallaria, Actinocoryne contractilis*, brown lines), and they are all characterized by pre-stored energy, either in the form of osmotic pressure or elastic energy, being released on a fast time scale^60^. The elastic jumpers like *Pseudotontonia* and *Tontonia* (purple lines) are right at the border of cAc ∼ 1, while hydrodynamic jumpers (green lines) and surface-based motility (blue lines) all have observed accelerations weaker than the naive hydrodynamic effect.

#### Ultrafast velocity

We next want to develop a quantitative framework to evaluate extreme velocity at the cellular scale. One convenient aspect of comparing velocities is that they can already be compared on the dimensionless scale of transient Reynolds number (Re_T_), the Reynolds number calculated using maximum velocity (Fig. 2B). We can see that although the majority of examples of fast speed at cellular scale still fall under low Reynolds number regime, a considerable number of examples are actually in the intermediate Reynolds number regime (1 *<* Re_T_ *<* 1000). For *Spirostomum*, the inertial effect at intermediate Reynolds number regime is actually critical for them to use the pressure pulse generated from their ultrafast contraction for long-range intercellular communication, which would not be possible at the low Reynolds number regime^32^. The inertial effect at intermediate Reynolds number regime is also critical for the success of *Zoothamnium pelagicum* (Fig. 1, #**18**) in moving at high speed using medusa-like motion.

Nonetheless, although transient Reynolds number is useful for determining flow characteristics, it is hard to predict the fundamental limit on speed from it. We thus need to first determine the expected velocity of an organism given its size.

##### Expected velocity at cellular scale

Interestingly, it has been previously shown that the expected maximum velocity for a given organism is about 10 BL/sec.^57^ Meyer-Vernet and Rospars argued that there are three numbers that are roughly constant across organisms, from bacteria to whales: mass density (*ρ ∼* 10^3^ kg/m^3^), applied force per cross-sectional area (*σ ∼* 200kPa)^61^, and the maximum power per unit mass (*b*_M_ *∼* 2kW/kg)^61^. Now consider an organism with characteristic length scale *L*, cross-section area *S*, and a striding frequency *f* . Its mass scales with *M ∼ ρSL*, its step size scales with *L*, and its force scales with *σS*. Thus the energy required per stride per unit mass *W* can be written as 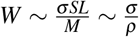 . Multiplying this by the striding frequency, we can derive the energy consumption rate per mass as *f W*, which should be limited by *b*_M_. The inequality *f W ≤ b*_M_ gives us the expected maximum normalized velocity as 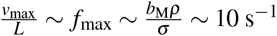. Meyer-Vernet and Rospars also showed that for a wide range of organisms, their maximum velocity usually falls between 1 BL/sec and 100 BL/sec.

However, one may question how much the above argument still holds for single cell organisms, many of which move around through cilia and flagella. Fortunately, similar conclusions can be derived from optimizing swimming efficiency and the hydrodynamics of cilia and flagella. Using Lorentz reciprocal theorem one can derive that the swimming velocity *v* of a microswimmer with body radius *r*, cilia length *𝓁*, averaged tangential stress at the surface *f*_*θ*_ swimming in a fluid of dynamic viscosity *μ* as 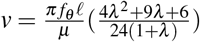, where 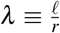. From the time-averaged swimming speed 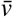, time-averaged drag force 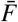, and the time-averaged mechanical output 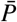, one can define the swimming efficiency of the microorganism to be 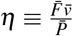.^62^ If the organisms do not change the number of cilia or flagella when they change sizes, the swimming efficiency will scale as *r*^*−*3^. This is because given a fixed number of cilia, increasing the radius of the organism would lead to *f*_*θ*_ ∝ *r*^*−*2^, *v* ∝ *r*^*−*2^, and *η* ∝ *rv*^2^ ∝ *r*^*−*3^.^62^ Therefore, microorganisms adjust their cilia or flagella density to ensure optimal efficiency. Based on the simulation performed by Omori et al., the optimal swimming speed of microswimmers are 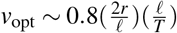, where *T* is the time period of ciliary beating. Substituting the usual ciliary beating frequency of 15.9 Hz^63^, we can show that the optimal swimming speed for microswimmers to be *v*_opt_ ∼ 25*r/*sec ~ 12BL*/*sec, similar to the results from Meyer-Vernet and Rospars.

Next, we collected the maximum speeds of many single-cell organisms with different behavioral phenotypes. The data was largely based on previously reported data^13,62,64,65^, with corrections and additional inclusion of ultrafast organisms (Fig. 2C). It can be seen that for most single-cell organisms, their maximum velocity indeed falls around 10 BL/sec. This again supports the definition of using 100 BL/sec as the cutoff for ultrafast velocity at the cellular scale. Many examples of them can be found in Figure S2.

##### Near linear scaling between cAc and Re_T_ suggests a universal energy conversion efficiency

Next, we reasoned that as both acceleration and velocity are ultimately limited by dissipation from the environment, there should be a scaling relationship between the two. Indeed, when we plot the transient Reynolds number with respect to the cellular acceleration number for various ultrafast cellular phenomena, we found a near linear scaling between the two, with a power law exponent of 0.945 (Fig. 2D), and this scaling holds across 6 orders of magnitude for both dimensionless numbers. This comes as a surprise, as cAc only contains information about acceleration but not velocity, while Re_T_ only contains information about velocity but not acceleration. However, as we revisit the definition of the two, we can show that 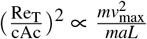, where the numerator is proportional to the maximum kinetic energy of the organisms, while the denominator is proportional to the work done across the length scale of the organism. A near linear slope between the two thus suggests a universal energy conversion efficiency for ultrafast cellular phenomena.

To calculate this energy conversion efficiency, we assume Re_T_ ≈ 5.21 cAc based on Figure 2D. Taking the square of both sides of the equation yields 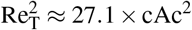. Yet from the definition of Re_T_ and cAc, we can show that 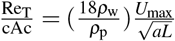.Since for most biological tissue, *ρ* ≈ *ρ*, taking the square of the equation we can show that 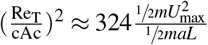. Comparing the two equations, we can show that 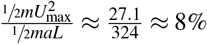. We will later show in Figure 5 how this energy conversion efficiency is useful for us to derive the fundamental limits on velocity and acceleration for several different mechanisms.

#### Ultrafast area strain rate

While the majority of the ultrafast mechanisms are associated with speed and acceleration, cells can also employ extreme changes in their volume, surface area, and density to achieve useful functions. In terms of surface area changes, most animal cell types change their cell surface area through an actomyosin-based mechanism.^49^ From the speed of a single motor head and the averaged length of a contraction unit, it can be shown that when the system is contracting at full speed with no external load, the maximum area strain rate would be 250%/sec.^49^ We can thus reasonably define any area strain rate greater than that as an ultrafast area strain rate.

Two examples of ultrafast area strain rate are known to date. (Fig. 3A) One is the ultrafast neck extension of *Lacrymaria olor*, where the area strain rate can go up to 400%/sec.^50^ The fast extension and retraction of the neck is critical for their effective search through space for prey,^47^ and they achieve this through a curve-creased origami mechanism to rapidly deploy the stored membrane reservoir.^50^ Another example is the ring cells of the constricting rings in nematophagous fungus. As the ring cell expands in volume to choke the passing nematode, the surface area also needs to change at a strain rate of 500%/sec^55^. This rapid expansion of surface area is possible because of the pre-synthesized membrane reservoir arranged underneath the cell membrane in a palisade shape, ready to be deployed.^55^ These two examples both highlight the power of quantitative analysis in identifying extreme cases that require novel mechanisms that cannot be explained through a traditional, actomyosin-based mechanism.

**Figure 3.**
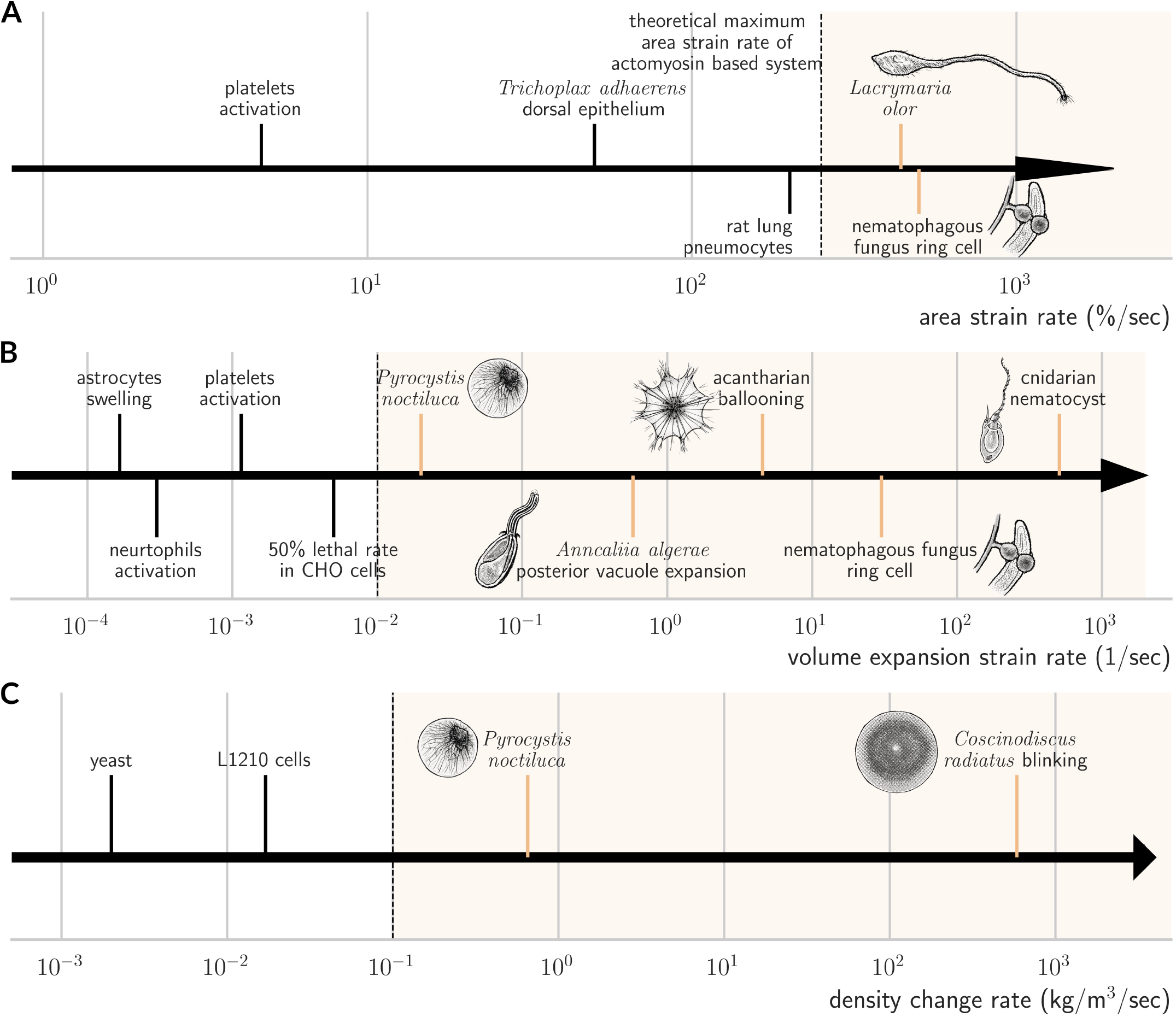
Comparison of ultrafast area, volume, and density change rates among various single-cell organisms.(A) Fast area strain rate at cellular scales. Previous studies showed that there is a maximal area strain rate limit of 250%/sec for actomyosin-based area changes^49^. We thus define any area strain rate beyond that limit as ultrafast. *Lacrymaria olor* utilizes curve crease origami to store their cell membrane for fast deployment^50^, while the ring cells of nematophagous fungus possess palisade-shaped membrane reservoirs to accommodate the rapid area changes during cell expansion^55^. (B) Fast volume expansion strain rate at cellular scales. We define volume expansion strain rate much greater than the 50% lethal rate in CHO cells during osmotic stress as ultrafast. The ultrafast volume expansion either happens in a separate compartment from the true cytoplasm (*Pyrocystis noctiluca, Anncaliia algerae*, acantharian) or is a single shot event (*Anncaliia algerae*, nematophagous fungus ring cells, cnidarian nematocysts). (C) Fast density change rate at cellular scales. We define density change rate much greater than the normal density change that happens during the cell cycle as ultrafast density change rate. *Pyrocystis noctiluca* accomplishes this through water influx and volume expansion in the central vacuole. *Coscinodiscus* accomplishes this through rapid exchange of higher density hydrated K^+^ ions and lower density hydrated Na^+^ ions in the cytosol.^54^ cnidarian species *Nematostella vectensis*, they carry a specialized voltage-gated calcium channel (nCa_*V*_) with voltage-dependent inactivation that can avoid triggering by non-prey water disturbance. When prey-derived chemical signals are detected by the sensory neuron, acetylcholine is synaptically released to the innervating nematocyte, causing hyperpolarization that relieves the inactivation of nCa_*V*_ channels, allowing for the mechanically triggered predatory attack.^80^ These examples highlight the fact that organisms evolve trigger logic according to the functional needs and the associated cost of executing ultrafast mechanisms.

#### Ultrafast volume expansion strain rate

In terms of volume expansion, several specialized mammalian cell types are known to expand their volume upon activation, such as astrocytes^66^, neutrophils^67^, and platelets^68^. However, their volume expansion strain rates are all around or below the order of 10^*−*3^/sec, as volume expansion beyond ∼5 × 10^*−*3^/sec will disrupt the biochemistry too much and cause cell death.^56^ We thus define any volume expansion strain rate greater than 10^*−*2^/sec (or 1%/sec) as ultrafast volume expansion. Among the identified ultrafast examples (Fig. 3B), they are either single-shot events (cnidarian nematocyst, nematophagous ring cells, and *Anncaliia algerae* posterior vacuole expansion) or have internal compartmentalization that separates the vacuole that is expanding from the true cytoplasm (*Pyrocystis noctiluca, Anncaliia algerae*, and acantharian). For *Pyrocystis noctiluca*, it even evolves a reticulated cytoplasm geometry wrapping around the large central vacuole, such that as the volume expands, the amount of membrane required to support the cytoplasm would only scale linearly with the size of the cell rather than quadratically.^51^ Both of the above examples, although different from translating in the fluid, still have to account for hydrodynamic dissipation for being embedded in the fluid. However, how much of that limits their strain rate is unclear so far.

#### Ultrafast density change

Finally, there are a few examples of organisms that can change their cell density at very fast speed to control their buoyancy. Most cells experience periodic density changes during their cell cycle as they swell up in volume before mitosis^69,70^. Nonetheless, the density change rate is usually on the order of 10^*−*2^ kg/m^3^/sec or less. We thus define any density change rate faster than 10^*−*1^ kg/m^3^/sec as an ultrafast density change rate (Fig. 3C). For *Pyrocystis noctiluca*, they can expand the volume by five times in 10-15 minutes, achieving a density change rate of 0.65 kg/m^3^/sec to avoid sinking to the bottom of the ocean.^51^ As for *Coscinodiscus*, they can rapidly change their density at a speed of 584 kg/m^3^/sec to achieve fast sinking, replace surrounding fluid, and optimize nutrient intake, highlighting the diverse range of functions that ultrafast cellular phenomena can accomplish.^53^

### Trigger mechanisms of cellular ultrafast motility

After establishing the quantitative framework for evaluating ultrafast cellular phenomena, we next explore the diverse range of mechanisms utilized by single cells to initiate ultrafast motility. Three broad classes of triggering mechanisms have been observed before, including chemically-triggered, mechanically-triggered, and chemically-gated mechanically-triggered (Fig. 4A). Examples of chemically-triggered ultrafast systems include microsporidia polar tube firing^71^ and the ultrafast swimming velocity observed in microaerophilic bacteria, magneto-aerotactic bacteria, and thermophilic hydrogenotrophic methanogens^11–13^. For microsporidia, as the spores need to germinate at appropriate environment (usually in the digestive system of the hosts), they are expected to evolve chemically-triggered mechanisms that sense specific pH levels, ionic compositions, or digestive products like sucrose.^71^ While various triggers for different microsporidia species have been identified, it remains unclear which proteins or receptors are activated by these triggers, and how this activation causes spore germination. (See Huang et al. 2023 for a review on this topic.^72^) For microaerophilic and magneto-aerotactic bacteria, the mechanism by which they sense oxygen gradients is still unclear, but it is likely related to the methyl-accepting proteins (MCP) family^73^. In *E. coli*, known oxygen sensors include the oxygen transducer Aer and Tsr protein, and their homologs can be found in the magneto-aerotactic bacteria^73^. For thermophilic hydrogenotrophic methanogens like *Methanococcus*, taxis towards hydrogen gas (or hydrogenotaxis) has also been experimentally demonstrated.^74^ The ultrafast speed in these cases allows them to rapidly move towards favorable niches in a steep gradient.

**Figure 4.**
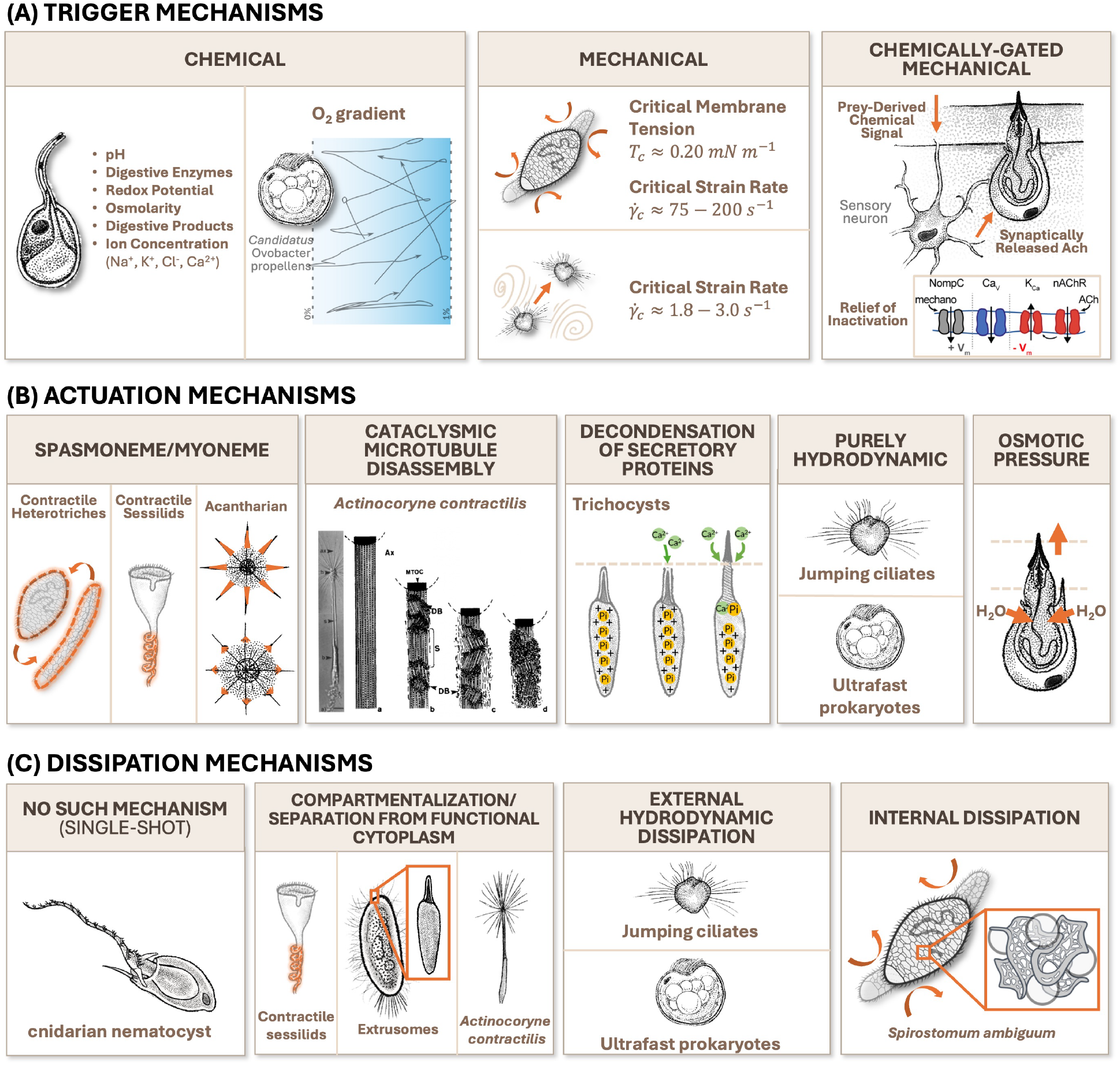
Ranges of trigger, actuation, and dissipation mechanisms in ultrafast cells. (A) We define trigger as the signal that initiates the ultrafast mechanisms. The trigger mechanisms of ultrafast cellular phenomena are either chemically-triggered (e.g., microsporidia^81^, ultrafast prokaryotes sensing oxygen gradient^11^), mechanically triggered (e.g., contractile heterotriches^32^, jumping ciliates^77^), or chemically-gated mechanical triggers (e.g., cnidarian nematocysts^80^). Purely mechanically triggered mechanisms are usually repeatable, with measurable critical strain rate 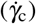 or membrane tension (*T*_c_). (B) Many ultrafast cellular phenomena actuate through releasing the energy stored in spasmonemes or myonemes upon Ca^2+^ signals (e.g., contractile heterotriches^82^, contractile sessilids^83^, and acantharians^84^). *Actinocoryne contractilis* performs stalk contraction through cataclysmic microtubule disassembly^38^, characterized by intercalary destabilization with intact microtubule arrays between destabilization bands, different from catastrophic microtubule depolymerization. *Paramecium* trichocysts are held in place by dissolved phosphate screening the positive charges in trichocyst matrix proteins. The calcium influx from the surrounding environment precipitates the dissolved phosphate, and trichocyst matrix proteins rapidly decondense due to charge repulsion^85^. The actuation mechanisms of jumping ciliates and ultrafast prokaryotes are purely hydrodynamics, characterized by long cilia or flagella with high beating frequency. (See Fig. 6.) Cnidarian nematocyst discharge is driven by osmotic pressure^9,86^. (See Figure 5 for detailed calculations.) (C) Cells with single-shot ultrafast motility do not need to evolve specific dissipation mechanisms. Organisms with contractile stalks (e.g., contractile sessilids and *Actinocoryne contractilis*) or extrusomes separate the ultrafast actuator from their functional cytoplasm to avoid damage. Most of the energy of ultrafast motility dissipates through external hydrodynamic drag, while *Spirostomum ambiguum* evolves an entangled architecture of rough endoplasmic reticulum and vacuolar meshwork to have an internal damping mechanism^59^.

**Figure 5.**
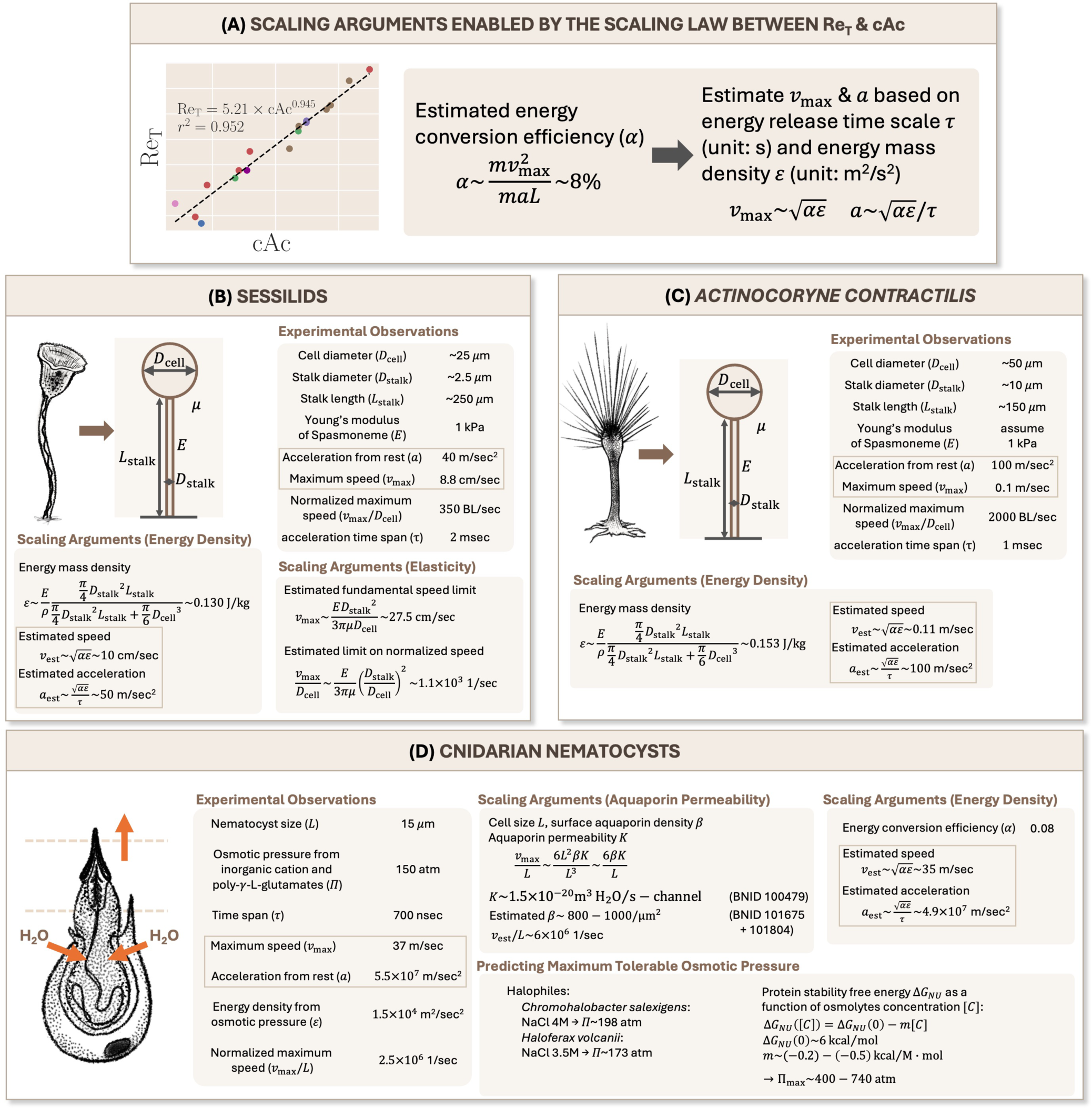
Test use cases for utilization of energy conversion efficiency scaling in ultrafast processes. (A) The scaling law between Re_T_ and cAc allows us to estimate an energy conversion efficiency (*α*) of 8% for ultrafast motility across 6 orders of magnitude and multiple behavioral phenotypes. We can estimate the fundamental limits on speed and acceleration if we can determine the energy mass density *ε* and the time scale of energy release *τ*. (B) Fundamental speed limits of sessilids. We model a contractile sessilid as a ball on a spring with Young’s modulus *E*, embedded in a viscous environment with viscosity *μ*. The actuation is based on the release of stored elastic energy in the spasmoneme in the stalk. A scaling argument based on the force balance between elastic forces and drag forces can provide an estimation of the speed limit but cannot predict the acceleration. Using the scaling argument listed in (A), we can estimate both the speed and acceleration of the contraction process. (C) Fundamental speed limits of *Actinocoryne contractilis*. Although we can simplify the morphology into a ball on a spring, the contraction of *Actinocoryne contractilis* stalks involves cataclysmic microtubule disassembly (see Fig. 4B). Thus, we cannot use the elastic force balance argument anymore. However, we can still estimate the speed and acceleration based on the estimated energy mass density. (D) Fundamental speed limit of cnidarian nematocysts. Cnidarian nematocyst discharge is driven by osmotic pressure, with key numbers summarized^9,86^. We can estimate the fundamental speed limit mechanistically based on the water permeability of aquaporins, but this does not allow us to estimate the acceleration. Using the scaling argument listed in (A), we can estimate both the speed and acceleration of an osmotic pressure-driven process. By comparing the osmotic pressure in nematocysts with the maximum tolerable osmotic pressure observed in halophiles and predicted from protein stability energy, we conclude that the osmotic pressure stored in nematocysts is approaching the theoretical limit for biological tissues.

Some other organisms rely primarily on purely mechanical triggers, and these are usually repeatable ultrafast mechanisms (in contrast to single-shot events). As the trigger mechanisms need to be implemented by the molecular components, the actuation time scale of the trigger mechanism for a repeatable process would set the minimal duration of the actuation time scale, so one trigger would not initiate multiple actuation. For purely mechanical trigger with repeatable deployment, the trigger is usually suspected to be some mechanical sensitive ion channels like Piezo proteins, and their actuation time scale is about 1-3 msec^75^. This is consistent with the observation that most repeatable, purely mechanical trigger ultrafast motility have an actuation time scale longer than millisecond range (Fig. S4). The sensitivity of the molecular components also sets the critical strain rate for trigger, which can be measured using a radial-flow microfluidics setup with a strain rate gradient along the streamlines. For *Spirostomum ambiguum*, the critical strain rate needed to trigger the ultrafast contraction has been shown to be around 75-200 sec^*−*1^, depending on the length of the cell^32^. Using regularized Stokeslet calculation, the corresponding membrane tension caused by the surrounding fluid strain is estimated to be 0.2 mN/m^32^. For *Stentor coeruleus*, although the absolute threshold in terms of critical strain rate has not been measured, mechanical sensitivity is clearly demonstrated in this organism, and habituation to mechanical stimuli has also been observed^33,76^. For several species of jumping ciliates and flagellates, the critical strain rates for their escape responses have been measured: 1.8 s^*−*1^ for *Balanion comatum*^77^, 2.4 s^*−*1^ for *Mesodinium pulex*^77^, 3.6 s^*−*1^ for *Strobilidium*^78^, 8.9 s^*−*1^ for *Chrysochromulina simplex*^78^, and 12.3 s^*−*1^ for *Gymnodinium*^78^. The critical fluid velocity that can trigger the ambush attack of *Mesodinium pulex* has also been measured to be between 15-42 *μ*m/sec, presumably sensed by the deformation of equatorial cirri^79^.

For single-shot ultrafast mechanisms, such as the firing of nematocysts in cnidarian, it has been shown that these mechanisms are mediated by a chemically-gated mechanical trigger to avoid futile firing caused by non-prey hydrodynamic signals.^80^ In cnidarian species *Nematostella vectensis*, they carry a specialized voltage-gated calcium channel (nCa_*V*_) with voltage-dependent inactivation that can avoid triggering by non-prey water disturbance. When prey-derived chemical signals are detected by the sensory neuron, acetylcholine is synaptically released to the innervating nematocyte, causing hyperpolarization that relieves the inactivation of nCa_*V*_ channels, allowing for the mechanically triggered predatory attack.^80^ These examples highlight the fact that organisms evolve trigger logic according to the functional needs and the associated cost of executing ultrafast mechanisms.

### Actuation mechanisms and energy source of cellular ultrafast motility

After considering a range of trigger mechanisms for ultrafast motility, we next examine a broad range of actuation mechanisms (Fig. 4B). Previous reviews of these supramolecular “springs and ratchets” broadly describe the controlled energy release in some of these actuation mechanisms^87,88^. Here we provide a detailed classification of these unique mechanisms. Furthermore, by applying the previously described scaling law, we re-evaluate the speed and acceleration limit predictions for selected examples.

#### Calcium-sensitive ATP-independent non-actin filaments and their fundamental limits

Many ultrafast motility are actuated by calcium-sensitive, ATP-independent non-actin filaments (often referred to as spasmonemes or myonemes), as seen in contractile heterotriches (*Spirostomum*^82^ and *Stentor*^89^) and contractile sessilids (*Vorticella, Zoothamnium*, and *Carchesium*)^34^. Experiments on glycerinated *Vorticella* stalks indicate that spasmoneme or myoneme contraction is mediated by the electrostatic screening of the negative charges intrinsic to the spasmoneme/myoneme proteins, as the stalk shortening can also be elicited by other divalent and trivalent cations.^90^ A mechanochemical model has recently been developed to combine the effect of traveling calcium waves, elasticity, and viscous drag to explain the contraction kinematics observed in contractile heterotriches and sessilids.^60^

Aside from these two large groups, myoneme is also responsible for the ultrafast volume expansion observed in acantharian^91^, the piston action of *Erythropsidinium*^92^, and even the contraction of several leptodiscinae dinoflagellates species^93^. In acantharians, as the myoneme bundle bridging the periplasmic cortex and the apex of the spicule contracts, it expands the periplasm and presumably uses this to control buoyancy.^91^ Note that although the haptonema in haptophytes (such as *Chryrochromulina*, Fig. 1, #**21**) also appears to contract, it is actually a coiling process with no length shortening, and it is caused by a calcium-dependent conformational change of microtubule array.^29^

To theoretically estimate their speed and accelerations, we turn to the scaling law we discovered between Re_T_ and cAc. (Fig. 5A) The slope in the near-linear scaling between the two allows us to calculate an energy conversion efficiency *α* of 8%. If we assume that we know the energy mass density *ε* (in units of J/kg, or m^2^/sec^2^), the time scale of energy release *τ* (in seconds), and the energy conversion efficiency *α* (unitless), the velocity *v* should scale with 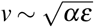, and the acceleration *a* should scale with 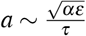.

For contractile sessilids like *Vorticella*, the system can be simplified to a spherical cell body with diameter *D*_cell_ attached to a contractile stalk with length *L*_stalk_, diameter *D*_stalk_, Young’s modulus *E*, and force constant *k*, embedded in surrounding fluid with dynamic viscosity *μ*.(Fig. 5B) The energy bearing part is only in the stalk, with a Young’s modulus of 1 kPa indicating an energy volume density of 1000 J/m^3^.^83^ Consider a *Vorticella* with *D*_cell_ = 25*μ*m, *L*_stalk_ = 250*μ*m, and *D*_stalk_ = 2.5*μ*m^94^, the energy mass density can be estimated as 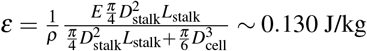, where *ρ* is the mass density approximated as 1000 kg/m^3^. For *Vorticella convallaria*, since the acceleration mainly occurs during the first 2 msec, we use this as our time scale *τ*. From this, we can estimate the velocity to be 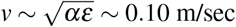, and the acceleration to be 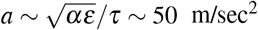. This estimation is extremely close to the experimentally observed maximum speed of 8.8 cm/sec and acceleration of 40 m/sec^2^.^94^

We can also estimate the fundamental limits of the system using the force balance between elastic forces and hydrodynamic drag (Fig. 5B). Assuming the hydrodynamic drag mainly comes from the cell body, the maximum velocity *v*_max_ can be estimated as

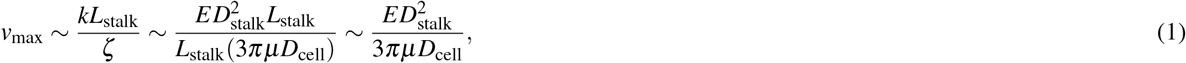

where *ζ* is the drag resistance of the cell body. If we normalize the maximum velocity with the size of the cell body, we get 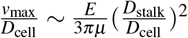. Given the Young’s modulus of spasmoneme has been previously estimated to be about 1 kPa^83^, and typically the stalk diameter is one-tenth of the diameter of the cell body, the theoretical limit of the maximum normalized velocity should be 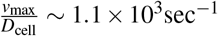, which is quite close to the experimental observation of 350 BL/sec.^94^

#### Cataclysmic microtubule disassembly and their fundamental limits

Actuation can also arise from rapid disassembly of filamentous structures in single cells. As seen in the rapid stalk and axopodial contraction in *Actinocoryne contractilis* and other heliozoan species (such as *Echinosphaerium*), disintegration of the filaments makes the recovery process more complex when compared to the contraction of non-actin filaments discussed before (Fig. 4B). Since the axopodia and stalk in heliozoa are composed of extensive microtubule arrays (which are arranged in a double-helical pattern in the axopodia of *Echinosphaerium*^95^ and a hexagonal pattern in the stalk of *Actinocoryne contractilis*^38^), their contractions also require the rapid breakdown of stalk and axopodial microtubules. Considering the extremely fast contraction speed (1-6 cm/sec for *Actinocoryne contractilis* stalk contraction^38^ and 0.6-1.5 mm/sec for *Echinosphaerium* axopodial contraction^96^), this cannot be explained by the catastrophic microtubule depolymerization, which has a shortening speed of approximately 35*μ*m/min.^97^ Electron microscopy studies later showed that the microtubule breakdown starts with a rapid intercalary destabilization, breaking the microtubule array into multiple destabilization bands sandwiching nearly intact microtubule arrays in between. As these destabilization bands expand, the microtubule segments tilt and continue to disintegrate into granular materials. Since this intercalary destabilization is a completely different disassembly process from the end depolymerization seen in catastrophic microtubule depolymerization, it is called ‘cataclysmic microtubule disassembly.’^38,98^ This intercalary destabilization might be caused by microtubule severing protein like katanin,^99^ though their causative role in heliozoa is yet to be determined.

For *Actinocoryne contractilis*, although morphologically it can also be simplified to a cell body attached to an energy-bearing stalk, similar to *Vorticella*, its much more complex mechanism prevents us from using the same elastic force balance arguments (Fig. 5C). However, as long as we can estimate the energy mass density, the scaling argument based on energy conversion efficiency and energy release time scale still allows us to predict speed and acceleration for *Actinocoryne contractilis* (Fig. 5C). Previous studies have shown that using the stress value of contractile sessilids can recapitulate the observed contraction forces in *Actinocoryne contractilis*.^36^ We thus assume the energy source in the stalk of *Actinocoryne contractilis* has similar mechanical properties and energy mass density as the spasmoneme. For an *Actinocoryne contractilis* with *D*_cell_ = 50*μ*m, *L*_stalk_ = 150*μ*m, *D*_stalk_= 10*μ*m^100^, the energy mass density can be estimated as 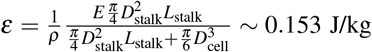. Using the same energy conversion efficiency of 8%, and since most of the acceleration occurs in the first 1 msec, we can estimate the velocity to be 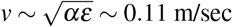, and the acceleration to be 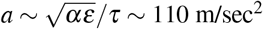. This estimation is again very close to the experimentally observed maximum speed of 0.1 m/sec and acceleration of 100 m/sec^2^.^100^

#### Decondensation of secretory proteins

In contrast to the rapid length shortening in the aforementioned examples, extrusomes like trichocysts are characterized by rapid length extensions (Fig. 4B). (See Plattner 2017 for a detailed review on *Paramecium* trichocysts.^44^) Structurally, a trichocyst is composed of a ∼3 *μ*m long and ∼ 0.3 *μ*m wide body part and a ∼ 2 *μ*m long tip part. The mature trichocyst (composed of trichocyst matrix proteins) has a cross-striated periodicity of 7 nm.^101^ When triggered (experimentally by secretagogue such as aminoethyldextran), it rapidly decondenses into 8 times its original length, and the cross-striation now has a periodicity of about 56 nm.^101^ The entire decondensation process is less than 1 msec, resulting in a speed greater than 24 mm/sec.^44^ During the rest state, the positive charges of trichocyst matrix proteins are screened by free phosphate ions dissolved within the trichocyst matrix. After the trigger and membrane fusion between trichocyst membrane and cell membrane, calcium influx from the surrounding precipitates the free phosphate ions, causing a rapid decondensation of the trichocyst matrix proteins^85^. The actuation mechanisms for other extrusomes, such as ejectisomes or toxicysts, are still under active research.

#### Osmotic pressure driven mechanisms and their fundamental limits

Some of the fastest known motility mechanisms in biology is claimed by other extrusomes, such as the nematocysts of cnidarian, that are driven by osmotic pressure (Fig. 4B, Fig. 5D).^86^ In cnidarians, 15 *μ*m large nematocysts are filled with poly-*γ*-glutamate and their associated cations, creating an osmotic pressure of 150 atm.^86^ The discharge happens on a time scale of 700 nsec, causing a speed of 37 m/sec (or 2.5 × 10^6^ BL/sec) and an acceleration of 5.4 × 10^6^*g*, exerting a pressure of 7.7 GPa at the site of impact.^9^ Using mercury chloride, a selective aquaporin inhibitor, it has also been demonstrated that the water transport of nematocysts are mediated through aquaporins.^102^ While the mechanistic and molecular details of cnidarian nematocyst discharge remain an active research area^103^, here we use scaling arguments to explore the fundamental limit of any osmotic pressure-driven mechanisms using biological tissues (Fig. 5D).

Mechanistically, as the ejection speed and volume expansion of nematocyst is fundamentally limited by water influx, knowing the size of the cell *L*, the surface aquaporin density *β* and the aquaporin permeability *K* allows us to estimate the maximum normalized speed as

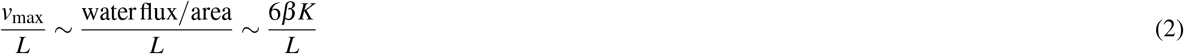

Even though the surface aquaporin density and aquaporin permeability are not known for cnidarian species, we can estimate them using data from other organisms. Based on data from human red blood cells (known for their fast water exchange capacity), we estimate the surface aquaporin density to be 800-1000 copies/*μ*m^2^ (using BNID 101675^104^ and BNID 101804^105^).

With some arithmetic, the water permeability of aquaporin can be estimated as *K* ∼ 1.5 × 10^*−*20^m^3^H_2_O*/*s − channel.^106^ Using these numbers, we estimate the normalized maximum velocity to be 6.6 × 10^6^ s^*−*1^, which is very close to the experimental observation of 2.5 × 10^6^ s^*−*1^ (Fig. 5D).

We can also apply our scaling argument based on energy mass density and the time scale of energy release to estimate the velocity and acceleration for cnidarian nematocysts (Fig. 5D). Since the energy here is stored in terms of osmotic pressure, we can estimate the energy mass density *ε* by dividing osmotic pressure Π by the mass density *ρ*, which gives 1.5 × 10^4^ m^2^*/*s^2^. Using the aforementioned time scale of 700 nsec and an energy conversion efficiency of 8%, we estimate the velocity to be 35 m/sec and the acceleration to be 4.9 × 10^6^*g*. This is again very close to the experimental observations of 37 m/sec velocity and 5.4 × 10^6^*g* acceleration,^9^ demonstrating the wide range of applicability of the energy conversion efficiency derived from the scaling between Re_T_ and cAc.

Finally, if osmotic pressure is so effective in generating ultrafast system, is it possible to further increase the osmotic pressure to engineer even faster biological systems? Interestingly, the osmotic pressure observed in cnidarian nematocysts seems to approach the theoretical limit that is tolerable by biological tissue (Fig. 5D). Halophile bacteria such as *Chromohalobacter salexigens* can maximally tolerate sodium chloride at 4M (Π ∼ 198atm),^107^ while another halophile bacterium *Haloferax volcanii* can tolerate salinity of 24% (Π ∼ 173atm).^108^ This maximum tolerable range can be predicted when we look at the protein stability in osmolytes. For most proteins, the stability energy (Δ*G*_*NU*_) falls in the range of −4 to −6 kcal/mol^109^, which is comparable to the energy released from ATP hydrolysis (-6.69 kcal/mol, BNID 100777^110^). In the presence of osmolytes with concentration [*C*], the stability energy of the protein can be written as Δ*G*_*NU*_ ([*C*]) = Δ*G*_*NU*_ (0) − *m*[*C*], where *m* is a constant which usually falls in the range between −0.2 ∼ −0.5 kcal/M-mol.^109^ If we take *m* = −0.35 kcal/M-mol and Δ*G*_*NU*_ (0) = −6 kcal/mol, the concentration [*C*^*\**^] when Δ*G*_*NU*_ ([*C*^*\**^]) = 0 is 17M, which corresponds to an osmotic pressure of 420 atm. As this is an estimation for an average protein, it is likely that some proteins more susceptible to osmotic pressure would denature before 420 atm. Based on these estimations, we can thus say that cnidarian nematocysts are potentially operating very close to the limit of osmotic pressure driven mechanism for biological tissues.

#### Pure hydrodynamic based mechanisms and their fundamental limits

Finally, organisms like jumping ciliates or ultrafast prokaryotes completely rely on hydrodynamics for their fast swimming (Fig. 4B). As we mentioned previously, the swimming velocity of a hydrodynamic swimmer in Stokes flow can be calculated as 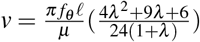 (Fig. 6C). Jumping ciliates, compared to other ciliates (statistics of their maximum speed shown in Figure 6A), thus carry several morphological and behavioral adaptations to increase their maximum swimming speed, such as increased ciliary beating frequency (increase *f*_*θ*_)^112^, longer cilia length (increase *𝓁*), fusion of cilia into membranelles, and longer cilia length relative to cell size (increase *λ*) (Fig. 6). Previous studies on *Halteria grandinella* showed that the cilia beating frequency during forward and backward swimming is about 105 Hz and 260 Hz, respectively, making them one of the systems with the highest cilia beating frequency.^112^ We also collected morphological data from many ciliate species and showed that indeed jumping ciliates tend to have longer cilia and a larger cilia length/half-cell length ratio (Fig. 6B-C). Note that even though in our model we did not explicitly account for the fact that the cilia in some of these organisms fuse into membranelle structures, the effect of cilia fusion is likely for cilia synchronization and its effect is included in the abstraction of cilia density into averaged tangential stress *f*_*θ*_ . Based on these statistics, we can predict that an averaged hydrodynamic jumper should achieve a velocity of 160 BL/sec, while a hydrodynamic jumper can maximally achieve 1060 BL/sec. Experimentally, the averaged normalized maximum velocity across different species is about 128 BL/sec, while the maximal record observed is about 361 BL/sec for *Strobilidium* sp.^16^

**Figure 6.**
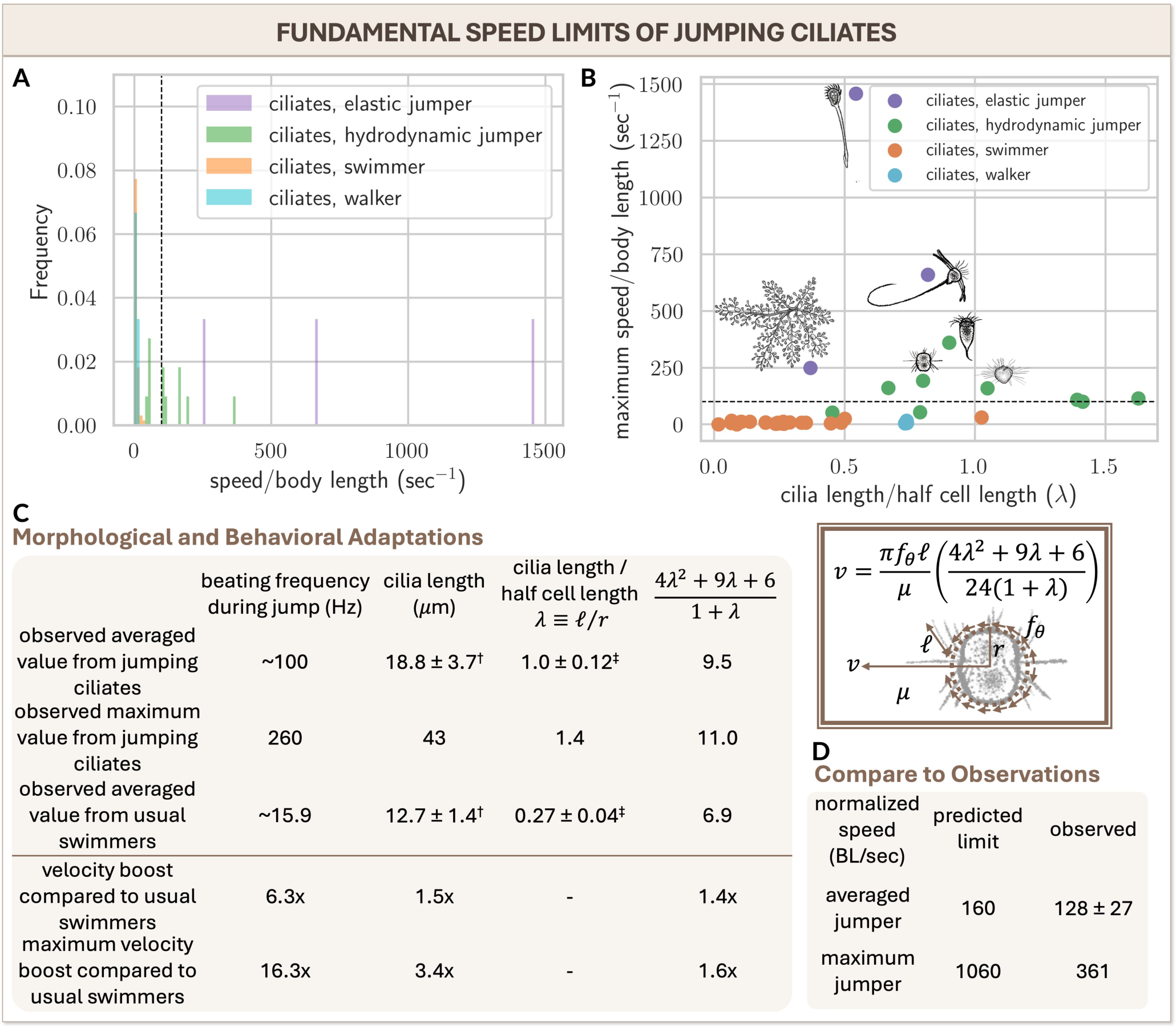
Scaling arguments on the fundamental speed limits for jumping ciliates. (A) Distribution of normalized maximum locomotion speed for ciliates with different modes of locomotion. The dotted vertical line indicates a velocity of 100 body lengths per second, the cutoff value we define as ultrafast velocity. Ciliates with ultrafast locomotion speeds are either elastic jumpers or hydrodynamic jumpers. (B) Hydrodynamic jumpers have longer cilia relative to their body length compared to hydrodynamic swimmers. Most hydrodynamic swimmers have a cilia length (*𝓁*) to half cell length (*r*) ratio (defined as *λ*) of less than 0.5, with *Strombidium sulcatum*^111^ as an exception. Walking ciliates (e.g., *Euplotes*) have long walking cirri but slow locomotion speeds. (C) Morphological and behavioral adaptations in jumping ciliates. The increased locomotion speed in jumping ciliates comes from their surge in cilia beating frequency and optimized cilia geometry. The term 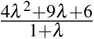 is a geometric factor that connects the tangential stress (*f*_*θ*_) generated by cilia to the predicted velocity (*v*) of a ciliate in media with viscosity *μ*, based on the Lorentz reciprocal theorem.^62^ Compared to a hydrodynamic swimmer, the averaged behavior of many jumping ciliate species can boost their velocity by 13.2 times, while the calculation based on the maximum observed across all jumping ciliates can boost their velocity by 88.7 times. (†: *p* = 0.038, one-way independent *t*-test. ‡: *p* = 2.6 × 10^*−*8^, one-way independent *t*-test.) (D) Comparison between the predicted and observed normalized speed (BL/sec) for the averaged behavior of jumping ciliates (denoted as “averaged jumper”) and the maximum capacity of jumping ciliates (denoted as “maximum jumper”). The predicted averaged normalized speed for jumping ciliates matches well with experimentally observed speeds. Currently observed fastest jumping ciliates have not reached the predicted maximum speed limit.

### Energy dissipation mechanisms of cellular ultrafast motility

After the cells receive the correct trigger and execute their ultrafast mechanisms, they next need to effectively decelerate themselves and dissipate the energy (Fig. 4C). This is critical, especially for single-cell organisms, as they need to remain functional and alive after the deployment of their ultrafast mechanisms. For some single-shot events like the ejection of cnidarian nematocysts, no such dissipation mechanism is required, as stem cells regenerate and replace the used nematocytes.^113^ For stalked protists like contractile sessilids, *Actinocoryne contractilis*, elastic jumpers, and extrusome-based mechanisms, the execution of ultrafast mechanisms is separated from their functional cytoplasm and thus has less impact on the internal organelle architectures. Most other behavioral phenotypes, such as hydrodynamic swimmers or hydrodynamic jumpers, mostly rely on external hydrodynamic drag to dissipate energy.

However, some ultrafast organisms, like *Spirostomum ambiguum*, utilize internal dissipation mechanism to decelerate themselves and dissipate energy.^59^ For *Spirostomum*, the contractile myoneme locates in the cortex and surrounds the functional cytoplasm. Based on the contraction kinematics, the extreme shape changes can generate a shear rate of 400 1/sec internally, which can potentially disrupt the organelle themselves or break their interorganelle connections. It turns out *Spirostomum ambiguum* has an entangled organelle architecture internally, where rough endoplasmic reticulum forms a fenestrated membrane sheet closely wrapping around the vacuolar meshwork throughout the entire cytoplasm.^59^ Simulations also show that the entangled organelle architecture can cause a strain-induced jamming transition inside the cell, which is shown to be the case inside the cell using spatial correlation of velocity directions of the vacuoles.^59^

Utilizing a jamming transition for energy dissipation enables the material to dissipate energy throughout its volume. Such an internal dissipation energy mechanism thus scales with the total volume of the cell, which is distinct from external hydrodynamic based dissipation mechanisms that can only scale with the surface area of the cell, making this a particularly appealing mechanism especially when cell size increases. So far, it is not clear how common this mechanism is for other contractile heterotriches or other ultrafast protists, but with the advances in fixation techniques and electron microscopy, we expect similar internal dissipation mechanisms to be more common than we currently think.

### Functions of cellular ultrafast motility

Finally, cells evolve with various ultrafast mechanisms to achieve specific functions that can benefit their survival (Fig. 7). Among all possible functions, predator escapes and predation are the obvious ones (Fig. 7A-B). In some cases, the ultrafast mechanisms are directly related to their success in predator escape. For example, hydrodynamic jumpers are experimentally demonstrated to be more effective in escaping from predators, especially filtration feeders (but less effective against ambush feeders).^77^ Cryptophytes such as *Rhodomonas* can also use the escape jumps generated by ejecting ejectisomes to escape from predators.^79^ Another interesting example is *Gymnodinium* against ambush feeder *Mesodinium* (which themselves are also hydrodynamic jumpers and can use their jump to escape as well). The fast swimming speed of *Gymnodinium* makes their approach feels like the approach of a predator and thus usually triggers the escape response of *Mesodinium* rather than their attack response, an elegant example of ‘hydrodynamic mimicry.’^79^ The defensive role of trichocysts is also experimentally demonstrated in *Paramecium* by comparing the survival rate of wild-type, trichocyst-deficient mutants, and artificially induced trichocyst-deficient cells in the presence of predators.^114^ Nonetheless, for some organisms, the ultrafast mechanisms only buy time for escape and by themselves are not effective if they lose their defensive toxins. For example, the ultrafast contraction of *Stentor* by itself does not prevent them from being eaten by *Coleps* if the contraction is not followed by the release of toxins from their cortical granules.^115^ The contraction gives them time and create distance to escape when the predator is paralyzed by the toxins, but it is not an effective escape strategy by itself.

**Figure 7.**
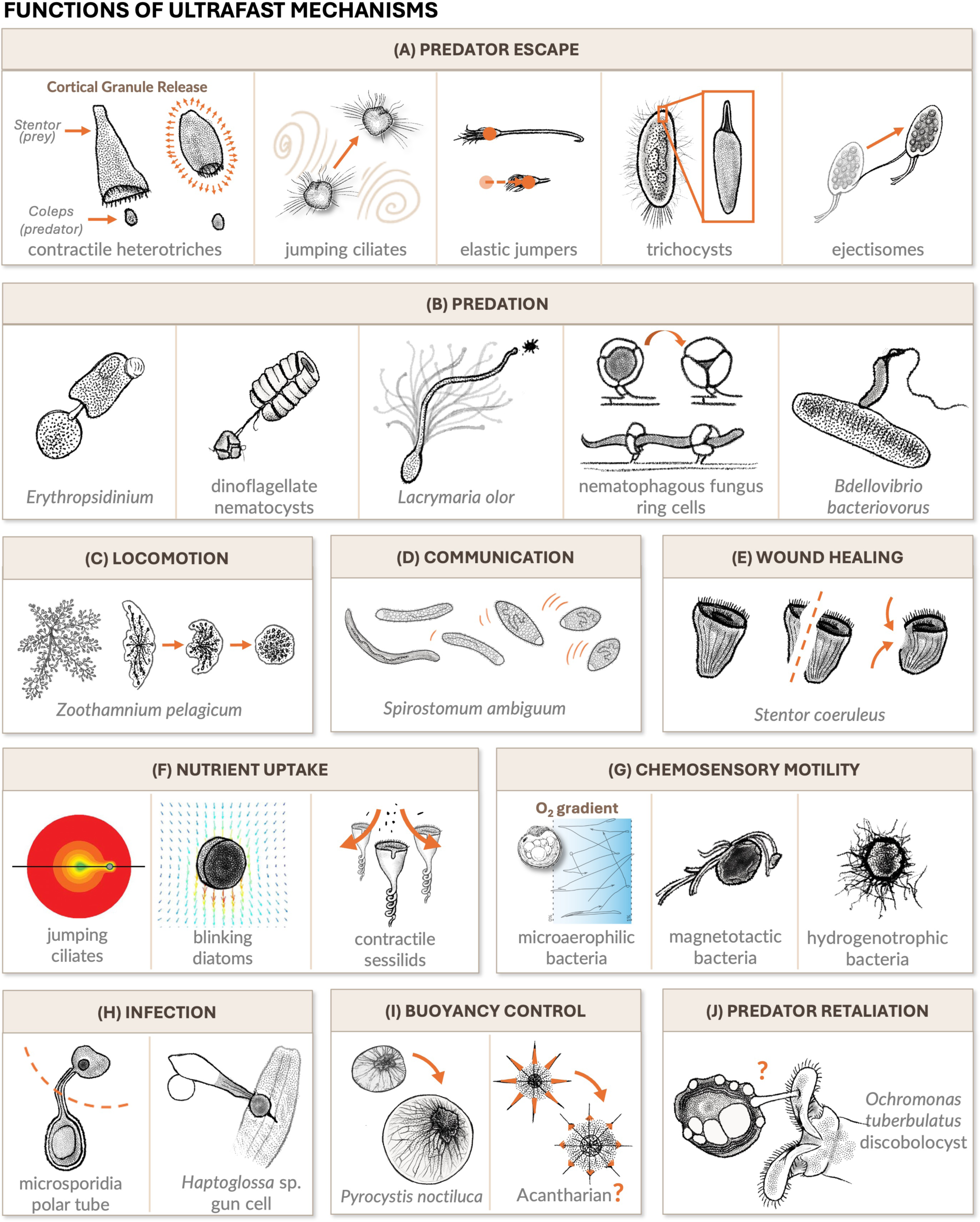
Cells utilize ultrafast motility for a diverse range of functions. Aside from the well-known functions in (A) predator escape, (B) predation, and (H) infection, they can also use them for (C) ultrafast locomotion^121^, (D) intercellular communication^32^, (E) wound healing^120^, (F) nutrient uptake^23,53,119^, and (I) buoyancy control^51^. (J) The discobolocysts of *Ochromonas tuberculatus* are hypothesized to have a role in predator retaliation as their high speed can potentially penetrate the cuticle of the predator, but this is not experimentally validated^43^. See Table S1 for the full list of functions and references.

As diverse as the predator escape responses in ultrafast cells are the predation strategies evolved in them (Fig. 7B). For example, *Erythropsidinium* can use the ultrafast extension and retraction of their piston to search for and ingest prey.^39^ They can also use their piston for fast locomotion (net speed ∼1 mm/sec, much faster than most dinoflagellates), although that happens much less frequently.^39^ Both cnidarian nematocysts and dinoflagellates nematocysts (in *Polykrikos* and *Nematodinium*) are used for predatory purposes. Note that although the two organelles share the same name, their phylogenetic origins are different.^116^ Several other offensive extrusomes are believed to be ultrafast as well, such as haptocysts, toxicysts and pexicysts (Table S1). However, as they are difficult to observe experimentally, no detailed kinematics have been reported. Some of these ultrafast players can also use their fast movement for more than one purposes. For example, for haptophytes such as *Chrysochromulina*, the rapid coiling of haptonema serves a triple role of prey capture^117^, locomotion by attaching to the surface^118^, and predator escape^29^.

Aside from these, ultrafast cells can also use their motility to optimize nutrient uptake (Fig. 7F). For example, some jumping ciliates’ fast jumps are not triggered by external fluid flow but instead can be spontaneous, as in the case of mixotrophic ones like *Mesodinium rubrum*.^23^ Using computational fluid dynamics studies, it has been shown that their fast jump can help them escape from their own nutrient-depleted boundary layer and expose them to water with fresh nutrients. Experimentally, it has also been demonstrated that the jumping distance is correlated to the pre-jump residence time, as a longer residence time requires them to jump farther away to get fresh nutrients.^23^ The rapid hydrodynamic jumps of *Mesodinium* thus have a triple role in predator escape, ambush feeding, and nutrient uptake. The fast contraction and slow extension of contractile sessilids are also shown to be an effective strategy to entrain nutrients.^119^ The fast swimming of *Candidatus* Ovobacter propellens can also increase the Sherwood number (defined as the ratio between convective mass transfer and diffusive mass transfer) of oxygen intake and is shown to give them 20% extra oxygen compared to being stationary.^11^ Similar to the case of *Candidatus* Ovobacter propellens, large diatom like *Coscinodiscus* can also use fast sinking to increase the Sherwood number of nutrient uptake to about 4.^53^

Another special function of ultrafast motility is its role in wound healing at cellular scale (Fig. 7E). Just like an animal’s connective tissues use wound contraction to reduce the size of the wound and assist in healing, *Stentor* also uses the contraction of the cortical myoneme to close its wounds. Through this, they can heal a 100 *μ*m-large wound in 100-1000 sec, achieving a wound healing rate which is at least 2 orders of magnitude faster than HeLa cells or fibroblasts.^120^

Some other functions, as we mentioned along the way, include the ultrafast locomotion of *Zoothamnium pelagicum*^121^ (Fig. 7C), the hydrodynamic intercellular communication of *Spirostomum ambiguum*^32^ (Fig. 7D), the infection of microsporidia^45^ and *Haptoglossa* species^46^ (Fig. 7H), and the buoyancy control as in the ultrafast volume expansion of *Pyrocystis noctiluca*^51^ and presumably also in acantharians^52^ (Fig. 7I). It is possible that cellular ultrafast players can use fast motility for other yet unknown functions. For example, it is hypothesized that *Ochromonas tuberculatus* can use the ejection of discobolocysts to attack the predator as retaliation due to the calculated high initial speed of 260 m/sec (Fig. 7J).^43^ However, this has not been experimentally validated. The full list of examples of known and unknown functions of cellular ultrafast motility can be found in Table S1.

## Discussion

In this work, we demonstrate that ultrafast motility is more widespread across the tree of life than previously believed. The survival benefits of these ultrafast mechanisms are evident from the diverse functions they serve. However, it remains unclear why microorganisms did not all engage in an evolutionary “arms race” of speed, or why certain evolutionary clades have not developed any ultrafast motility, whereas we see an overabundance of ultrafast examples in the SAR supergroup. Since evolution operates by repurposing, modifying, and building upon existing cellular and organismal frameworks — also known as “evolutionary tinkering”^122^ — future research should comprehensively consider the geometrical, structural, energetic, and ecological constraints in the evolution of novelty in biology.

By exploring extremes in biology and their fundamental limits, we may also uncover new molecular mechanisms yet to be discovered. For instance, the extreme area strain rates observed in *Lacrymaria olor* and nematophagous fungi suggest that traditional actomyosin-based mechanisms cannot account for their rapid area changes. This has prompted more detailed structural studies, which revealed the roles of curve-creased origami^50^ and pre-synthesized pallisade-shaped membrane reservoirs^55^ in enabling rapid membrane deployment. The quantitative framework proposed in this work can thus serve as a diagnostic tool in future research, helping to identify intriguing examples that may involve previously unknown molecular mechanisms.

In this work, we also present a near-linear scaling relationship between the cellular acceleration number (cAc) and the transient Reynolds number (Re_T_) for ultrafast cellular motility across six orders of magnitude. The slope of this scaling suggests an energy conversion efficiency of 8%, which we use to estimate the speed and acceleration limits of various mechanisms. Understanding these limits not only presents new biochemical and mechanical puzzles to be resolved but also has the potential to inspire and guide the design of soft robotics. For example, the spasmoneme contraction of *Vorticella* has already inspired the design of kirigami-based sensors and actuators^123,124^. It will be fascinating to see how the quantitative framework outlined in this work can predict and guide the development of future robotic devices.

Despite presenting the most comprehensive dataset of ultrafast cellular motility to date, with scaling laws derived from 17 acceleration data points across 2 phylogenetic domains and 4 phylogenetic supergroups, several limitations remain. Firstly, while our data points span the scale relatively evenly, acceleration data are generally difficult to obtain from the literature, as most studies report only velocity. Secondly, since the work covers examples ranging from low Reynolds number to intermediate and also a few transient high Reynolds number - it is possible that a more detailed framework might be needed to find universal fits for dissipation and total energy available in specific examples. At low Reynolds number examples, such as *Halteria grandinella*, the force balance between propulsion and drag is nearly instantaneous, observed acceleration is likely more related to the internal capacity to rapidly alter propulsion forces rather than to energy dissipation into the surroundings.

## Conclusion

In conclusion, we have provided a new systematic framework for studying ultrafast motility at the cellular scale and quantitatively evaluated them across the tree of life. We propose a new non-dimensional number and discover a new scaling relationship between cellular acceleration number and transient Reynolds number, which suggests a universal energy conversion efficiency of 8% for ultrafast cellular motility across six orders of magnitude. This energy conversion efficiency further allows us to estimate the speed and acceleration limit even when some specific details of the mechanism are not yet known. We also provided an extensive classification of known trigger mechanisms, actuation mechanisms, dissipation mechanisms, and functions of various ultrafast cellular motility. We believe this work lays the foundation to evaluate many yet-to-be-discovered ultrafast motility, and the quantitative framework we proposed can also give directions for the design of engineered biological systems or bio-mimetic soft robots.

## Acknowledgement

We thank all members of the Prakash Lab for scientific discussions and comments on figures. We thank Rebecca Konte for her tremendous help on figures. We thank Prof. Joshua Shaevitz and Rahul Chajwa for their insights on evaluating acceleration in low Reynolds number context. RC gratefully acknowledge support by Stanford University Bio-X SIGF Fellows Program and by the Ministry of Education in Taiwan. This work was support by HHMI Faculty Fellows Award (M.P), BioHub Investigator Fellowship (M.P), Keck Foundation Research Grant, Schmidt Foundation and NSF CCC (DBI1548297) and Gordon and Betty Moore Foundation.

## Code and Data Availability

All data and codes are available in Github (jrchang612/ultrafast_review).

## Materials and Methods

### Data collections

The data displayed in Figure 2A-B, D and Figure 3 was collected through a targeted search of known examples with ultrafast motility. Velocities, accelerations, area strain rates, volume expansion strain rates, or density change rates are either directly reported in the paper or calculated based on the reported data, with assistance from Fiji or Engauge Digitizer.

The data displayed in Figure 2C and Figure 6A was collected from Lisicki et al. 2019^64^, Herzog and Wirth 2012^13^, Nakamura 2020^65^, with additional the known examples of ultrafast organisms.

The morphological data displayed in Figure 6B was collected by direct measurements of microscopy images found in publications or plankton image database. The measurement was made using Fiji. See Section “Code and Data Availability” for the link to data.

### Statistical methods

The regression line and *R*^2^ in Figure 2D was calculated by simple linear regression on log(Re_T_) with respect to log(cAc).

The statistical testing on the morphology of hydrodynamic jumpers versus usual swimmers in Figure 6 was calculated using one-way independent *t*-test, assuming the hydrodynamic jumpers have a longer cilia length and larger *λ*.

### Reused figures

Part of Figures 4 and 7 contains reused figures from existing publications. The summary of their figure sources and copyright permission is in Table S2 in the Supplementary Material.

## Author contributions

R.C. and M.P. designed research; R.C. performed research; R.C. contributed new analytic tools; R.C. analyzed data; and R.C. and M.P. wrote the paper.

