## Supplementary Information for "Biophysical limits of ultrafast cellular motility"

### Biophysical limits of ultrafast cellular motility
#### Supplementary Information

**Ray Chang**[1] **and Manu Prakash**[1,2,*]

[1]Department of Bioengineering, Stanford University, Stanford, California, United States of America
[2]Woods Institute for the Environment, Stanford University, Stanford, California, United States of America
[*]

#### Contents

**1 Supplementary Figures** **2**
**2 Supplementary Tables** **6**
**References** **7**

### 1 Supplementary Figures

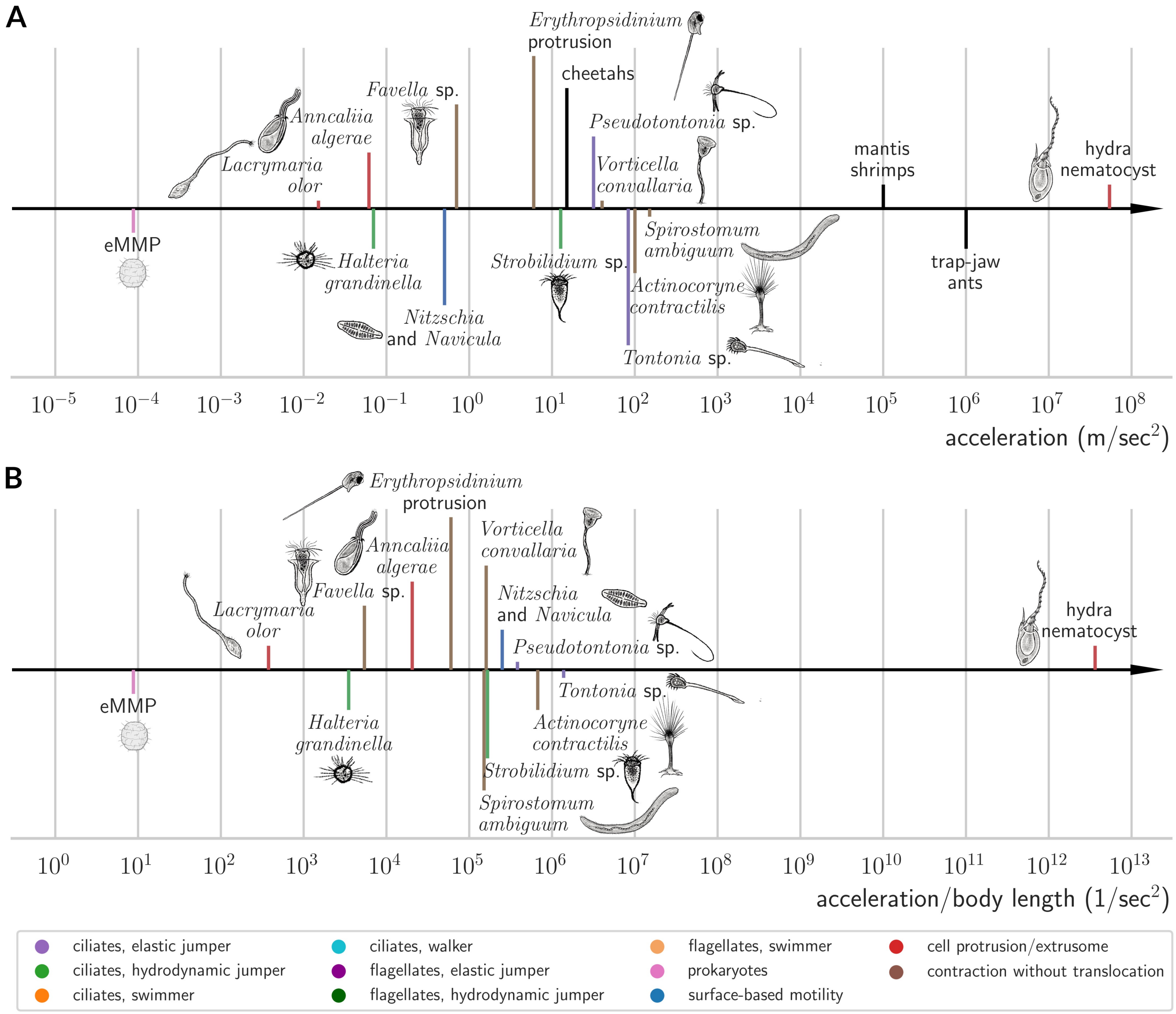


**Figure S1.** Fast accelerations at cellular scales. (A) A compilation of examples showing fast accelerations at cellular scales, with different colors representing various behavioral phenotypes. Black lines represent multicellular examples for reference. (B) Fast accelerations normalized by body length at cellular scales. Though being a common practice, this approach is not justifiable as discussed in detail in the main text.

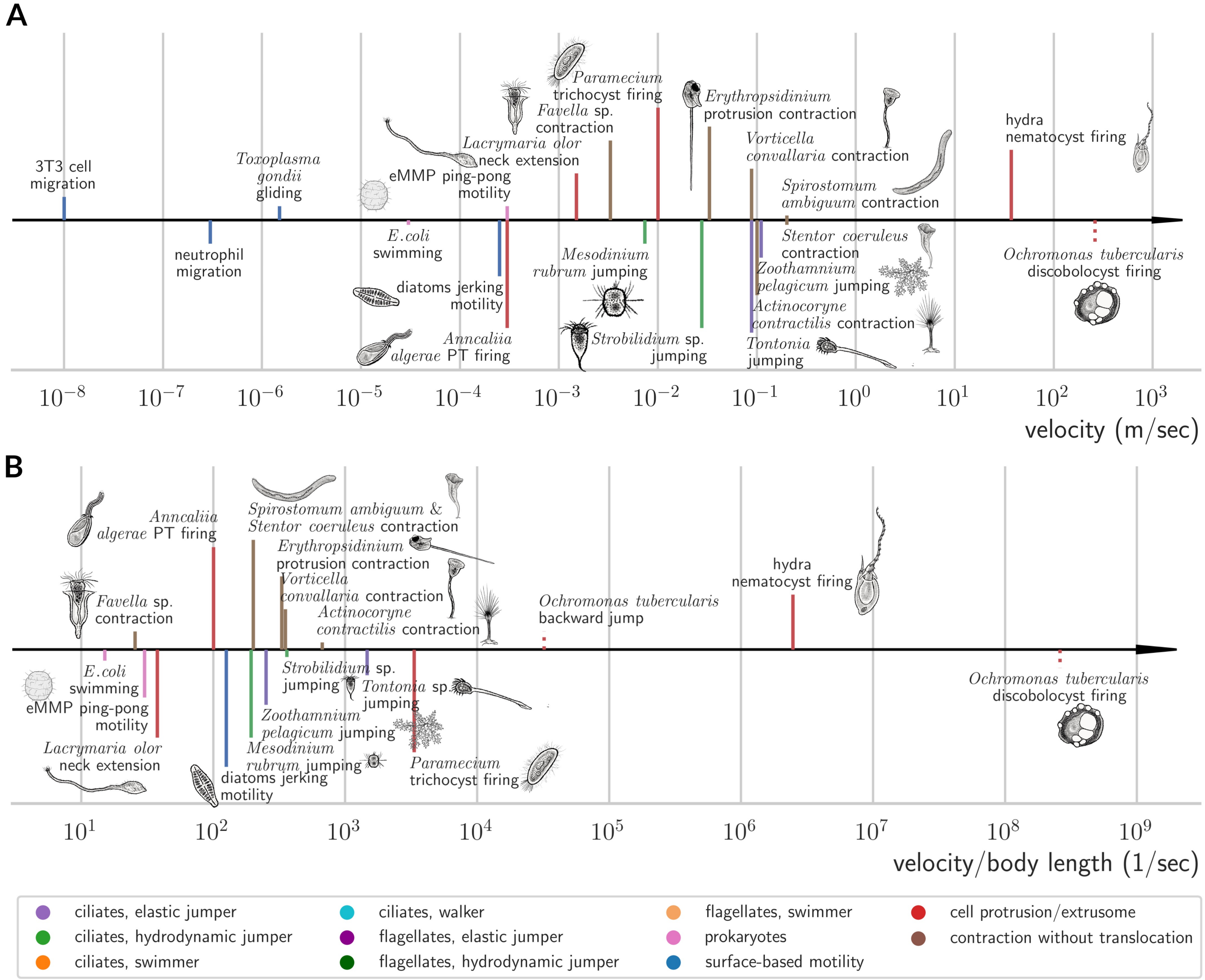


**Figure S2.** Fast velocities at cellular scales. (A) A compilation of examples showing fast velocity at cellular scales, with different colors representing various behavioral phenotypes. Dashed lines denote examples deduced by physics but lacking experimental confirmation. (B) Fast velocities normalized by body length at cellular scales.

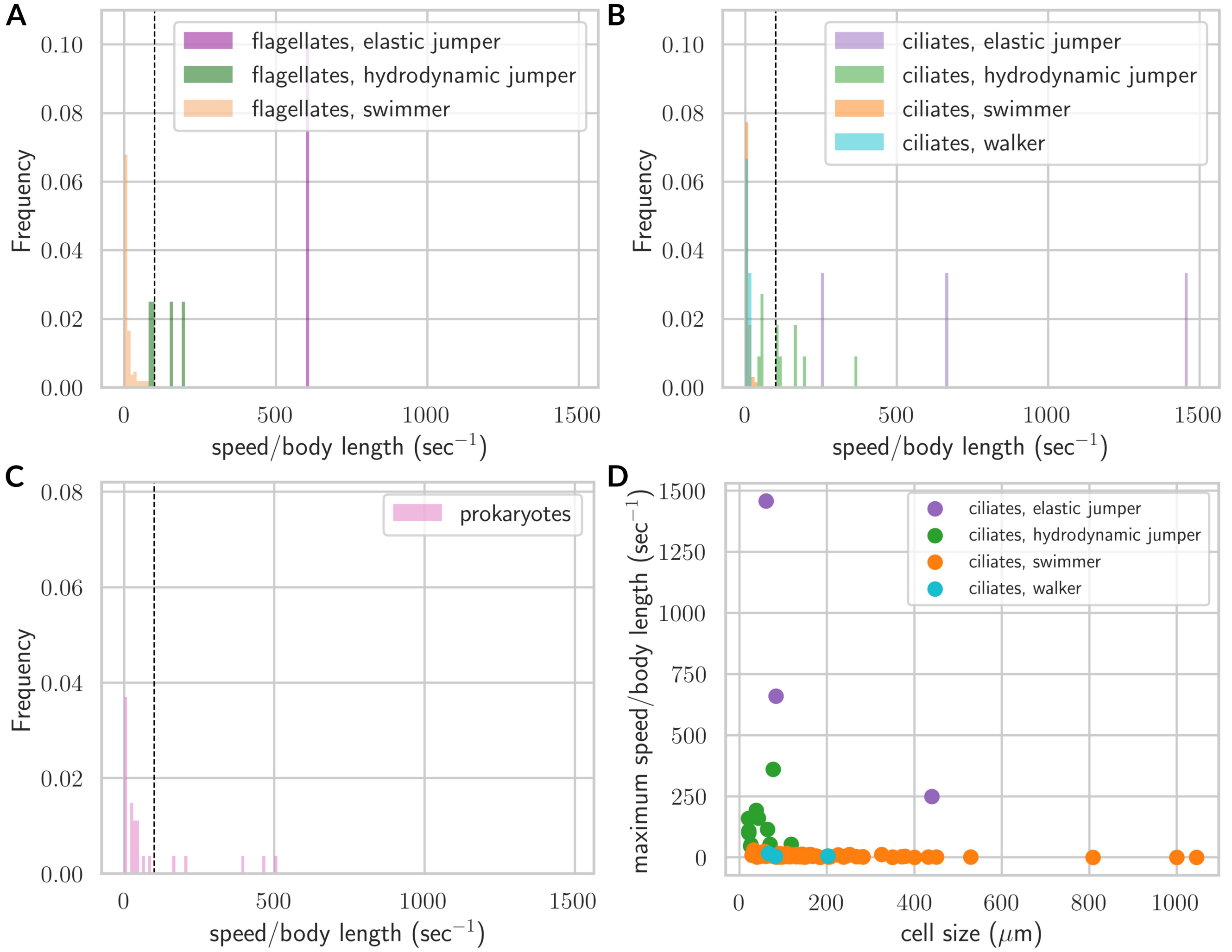


**Figure S3.** (A-C) Probability distribution of translocation speed in (A) flagellates, (B) ciliates, and (C) prokaryotes, grouped by behavioral phenotypes. (D) Maximum translocation speed normalized by body length with respect to cell size for ciliates. Note that most hydrodynamic jumpers have a cell size less than 100 $\mu$m.

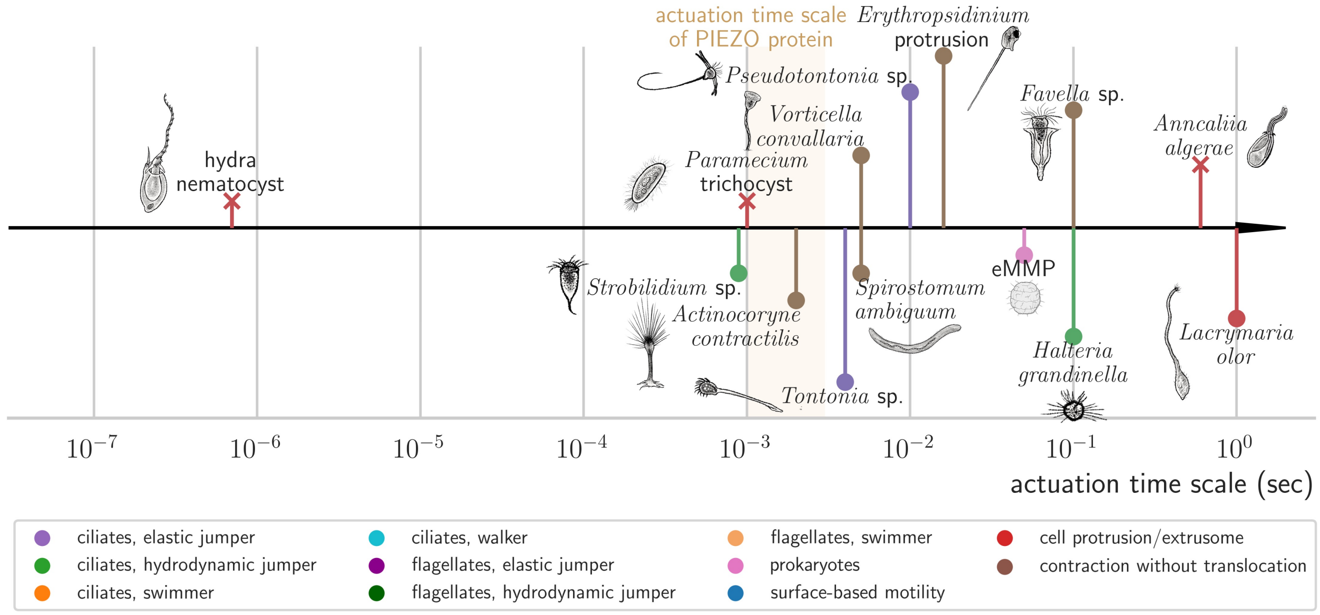


**Figure S4.** Actuation time scales for ultrafast cellular motility. A compilation of examples showing fast actuation time scales for ultrafast single-cell motility, with different colors representing various behavioral phenotypes. Marker 'o' indicates repeatable motility, while marker 'x' denotes single-shot deployment. The typical actuation time scale of PIEZO proteins (∼ 1-3 msec)[1] corresponds to the lower bound of most repeatable ultrafast motility.

#### 2 Supplementary Tables

**Table S1.** Functions of cellular ultrafast motility.

| Functions | Examples |
|---|---|
| escape response | contractile heterotrichs (*Spirostomum*[sp,ac][2], *Stentor*[sp,ac?][3]), contractile sessilidas(*Vorticella*[sp,ac], *Zoothamnium*[sp,ac?], *Carchesium*[sp,ac?])[4], *Actinocoryne contractilis* stalk contraction[sp,ac][5], elastic jumpers[sp,ac] (*Tontonia*[6], *Pseudotontonia*[6], *Spirotontonia*[?][7]), hydrodynamic jumping ciliates[sp] (*Strobilidium*[6], *Mesodinium*[8], *Uronychia*[9], *Halteria*[10], *Balanion*[11], *Askenasia*[?][12], *Cyclidium*[?][12], *Urotricha*[?][12]), hydrodynamic jumping flagellates[sp] (*Yihiella yeosuensis*[13], *Biecheleriopsis adriatica*[14], *Ansanella granifera*[15], *Bodo saltans*[?][16], *Resultomonas moestrupii*(formerly *Pedinomonas mikron*)[?][17], Heterocapsa rotundata[x][18], Gymnodinium simplex[x][19]), tintinnids contractions[x][20], ciliates trichocyst (*Paramecium*[sp][21,22], *Frontonia*[¿sp?], *Apofrontonia*[¿sp?], *Lembadion*[¿sp?], *Urocentrium*[¿sp?], Microthoracidae[¿sp?], *Swedmarchia arenicola*[¿sp?]), *Ochromonas tuberculatus* discobolocyst[sp†][23], *Strombidium* trichites[¿sp?][24], Leptodiscinae dinoflagellates[¿sp?,ac?] (*Cymbodinium elegans*, *Leptodiscus medusoides*, *Leptophyllus dasypus*)[25], cryptomonads ejectisomes[sp] (*Cryptomonas*, *Rhodomonas*, *Storeatula*)[26], *Chrysochromulina* haptonema coiling[sp][27] |
| locomotion | *Zoothamnium pelagicum*[sp,ac?][28], *Erythropsidinium*[x*][29], diatoms jerking[sp][30], *Sticholonche zanclea* axopod motion[sp?][31], *Chrysochromulina* haptonema coiling[sp][32] |
| communication | *Spirostomum ambiguum*[sp,ac][33] |
| predator retaliation¿ | *Ochromonas tuberculatus* discobolocyst[sp†][23] |
| nutrient uptake† | *Mesodinium rubrum*[sp][34], contractile sessilidas[sp,ac][35], *Stentor*[¿sp,ac?], *Coscinodiscus*[de][36], *Candidatus* Ovobacter propellens[sp][37] |
| chemosensory motility | *Candidatus* Ovobacter propellens[sp][37], *Methanocaldococcus jannaschii*[sp][38], *Methanocaldococcus villosus*[sp][38], *Magnetococcus marinus*[sp][39] |
| predation | *Erythropsidinium*[sp*][29], cnidaria nematocysts[sp,ac,vo][40], dinoflagellates (*Polykrikos*, *Nematodinium*) nematocysts[sp?,ac?][41], *Lacrymaria olor*[ar][42], *Didinium* pexicyst[?][43], suctorian haptocyst[?][24], Haptoria toxicyst[?][24], heliozoan axopod contraction[ac?,sp][5], *Bdellovibrio bacteriovorus*[sp][44], nematophagous fungus (*Drechslerella dactyloides*, *Arthrobotrys oligospora*, *Arthrobotrys brochopaga*) ring cell[vo,ar][45], hydrodynamic jumpers (*Mesodinium rubrum*[sp][46], *Yihiella yeosuensis*[sp][13], *Ansanella granifera*[sp][15], *Bodo saltans*[sp?][16]), *Chrysochromulina* haptonema coiling[sp][47] |
| infection | microsporidia polar tube[sp][48], *Haptoglossa mirabilis* gun cell[sp?][49] |
| wound healing | *Stentor coeruleus*[sp,ac?][50] |
| buoyancy control | *Pyrocystis noctiluca*[vo,de][51], acantharian ballooning[¿vo], *Coscinodiscus*[de] |
| unknown | contractile Dinophycean flagella[x][52], *Trachelonema* and *Tracheloraphis* rhabdocyst[?][53], *Remanella* nematocyst and orthonematocyst[?][54] |

Notation: sp: ultrafast speed; ac: ultrafast acceleration; vo: ultrafast volume expansion; ar: ultrafast area strain rate.; de: ultrafast density change.
?: likely ultrafast but lacking quantitative evidence.
†: theoretical evidence only.
¿: likely to have this function but not a consensus nor experimentally confirmed.
x: spring and latch mechanism but does not reach ultrafast.
*: For *Erythropsidinium*, the motility of the piston is an ultrafast speed phenomenon, but the locomotion of the cell body is not ultrafast.

**Table S2.** Summary of figure sources on reused figures.

| Figures | Sources | Copyright status |
|---|---|---|
| Figure 4A "Relief of inactivation" | Weir et al., 2020[55] Fig. 6A | Figure reuse permitted by CC BY 4.0. |
| Figure 4B "Cataclysmic microtubule disassembly" | Febvre-Chevalier and Febvre, 1992[56] Fig. 11 | Figure reuse permitted by CC BY-NC-SA 4.0. |
| Figure 4C "Internal dissipation" | Chang and Prakash, 2023[57] Fig. 1A, Fig. 2G | PNAS allows authors to use their original figures in their future works without needing to obtain permission. |
| Figure 7 "Nutrient Uptake" - jumping ciliates | Jiang and Johnson, 2017[34] Fig. 6 | Permission explicitly obtained from Copyright Clearance Center. |
